# Viral load-driven systemic immune exhaustion is an enabler of antibody breadth in HIV infection

**DOI:** 10.64898/2026.06.05.730519

**Authors:** Izumi de los Rios Kobara, Uma M Mangalanathan, Soneida Deline-Caballero, Kassandra Pinedo, Mikayla Stabile, Camilo Espinosa Bernal, Hannah L. Itell, Vrasha Chohan, R. Scott McClelland, Kishorchandra Mandaliya, Julie Overbaugh, Catherine A. Blish

## Abstract

A broadly neutralizing response to a human immunodeficiency virus (HIV) vaccine is a major goal of the field, yet the determinants of antibody breadth remain poorly defined. Antibody responses to HIV evolve over years of infection, developing unusual features such as high mutation rates in the subset of individuals that acquire breadth. Using single-cell RNA sequencing, single-cell proteomics, and plasma neutralization assays in a longitudinal Kenyan cohort of treatment-naive women living with HIV, we identify systems-level immune correlates of antibody breadth. Broad and narrow neutralizers diverge *early* in infection along a viral load-driven axis with broad neutralizers exhibiting greater viral loads and CD4 T cell decline; concurrently, NK cells, CD8 T cells, and monocytes, broad neutralizers displayed greater magnitude of immune activation in early infection, followed by exhaustion in late infection. This functional decline may dampen NK cell-mediated pruning of T follicular helper cells, creating a permissive environment for sustained germinal center activity and antibody maturation. Narrow neutralizers, by contrast, maintain functional cellular immunity but lack the antigenic pressure and permissive exhaustive state associated with breadth. Together, these findings suggest that antibody breadth in HIV infection reflects failed viral control rather than a successful antiviral response with implications for vaccination and cure strategies that must balance cellular and humoral immunity.

## Introduction

Human Immunodeficiency Virus (HIV) remains a critical global health crisis with 40.8 million people living with the virus and 1.4 million people infected in 2024^1^. There will be an estimated 73% increase in HIV infections over the next 5 years just in the United States due to health funding cuts^2^, emphasizing the continued need for a vaccine or cure. HIV primarily infects CD4 T cells, leading to a gradual decline in immune function and death from acquired immunodeficiency syndrome (AIDS) in untreated individuals^3–6^. A vaccine to prevent transmission is a key step in ending the HIV epidemic^7^. A rare subset of people living with HIV (PLWH) develop broadly neutralizing antibodies (bnAbs), antibodies with cross-tier, cross-strain neutralizing capacity^8,9^. BnAbs arise only after years of untreated HIV infection and display unusual features including extremely high somatic hypermutation (SHM, in some cases greater than 30%), long CDR3 sequences, and autoreactivity^8,10–14^. Passively transferred bnAbs can provide protection from infection in non-human primates and can suppress viral rebound in human clinical trials, providing strong rationale for their elicitation by vaccination^15–28^. Vaccine trials have thus far failed to produce bnAbs in healthy individuals; however, recent progress engineering germline-targeted immunogens offer considerable promise^29–31^. Several host and viral factors have been associated with bnAb development including high viral load, envelope protein diversity, elevated T follicular helper (Tfh) cells, and natural killer (NK) cell dysfunction^14,32,33^. How host and viral determinants interplay to enable antibody breadth and whether these features emerge in early infection remain unresolved. Understanding how systems-level immune dynamics translate antigen burden and immune activation into broad antibody responses is critical for the rational design of vaccines capable of eliciting bnAbs in healthy individuals.

Limited transcriptomic data exists for PLWH who develop bnAbs. To date, the only study to examine the transcriptome of individuals with bnAbs used bulk transcriptomic profiling followed by qPCR in sorted immune cell types, identifying NK cell dysfunction as the sole association with bnAbs^34^. NK cells are innate lymphocytes with anti-viral capacity, including direct killing of HIV infected cells^35–38^, but are underappreciated regulators of antibody responses^34,39–50^. In mice *in vivo* and in human *in vitro* co-cultures, NK cells can kill Tfh cells, limiting antibody responses^45,46,51^; NK cell activity is also correlated with poor response to vaccination in human trials^42–44^. We recently demonstrated in acute SARS-CoV-2 infection that NK cells from individuals with narrow antibody breadth expressed cytotoxic and interferon-stimulated gene (ISG) programs, and that ISG-driven NK cell activation suppressed B cell responses and killed Tfh-like cells *in vitro*^51^, suggesting that NK cell mediated suppression of adaptive immunity directly limits breadth. Finally, a small single-cell RNA-sequencing (scRNA-seq) study of hyperacute HIV infection found early cytotoxic, proliferating NK cells in the 2 out of 4 participants who displayed relative viral control although antibody breadth was not measured^52^. Chronic HIV infection demands far greater SHM and germinal center persistence than acute SARS-CoV-2 infection or vaccine responses. How specific NK cell states and their temporal evolution shape the divergence between broad and narrow antibody responses remains poorly studied and unresolved at both bulk expression and single-cell resolution.

Characterization of the immune response to untreated HIV infection at the single-cell level remains limited. ScRNA-seq studies of HIV have focused on small cohorts^53,54^, acute infection^52,55^, antiretroviral therapy (ART)-treated cohorts^56–60^, or specific cell subsets^61,62^, leaving the systems-level immune dynamics and longitudinal analysis across years of untreated infection, and their relationship to antibody breadth, largely unresolved. Here we address this gap by performing the largest single-cell characterization of untreated HIV infection to date in a longitudinal cohort of Kenyan PLWH, allowing us to capture the full disease course within the same individuals. At the time of sample collection ART was not available to PLWH in Kenya who had not yet progressed to AIDS, enabling a study design that can no longer be ethically replicated given that ART is now the standard of care at diagnosis globally. Our approach integrates 10X 5’ scRNA-seq with paired V(D)J B cell receptor (BCR) sequencing, mass cytometry by time of flight (CyTOF) for single cell proteomics, and plasma neutralization assays to understand systemic immunological determinants of antibody breadth. We recapitulate findings that breadth in late infection is correlated with higher viral load; however, we show for the first time that immunological trajectories diverge early in infection. Higher antigen burden in broad neutralizers drives systems-level immune activation that transitions to exhaustion and dysfunction in late infection. In NK cells, this is consistent with the only bulk expression study performed linking NK cell dysfunction to bnAb development in chronic HIV infection^34^. Narrow neutralizers retain cytotoxic NK cells in late infection, highlighting a phenotype consistent with Tfh cell pruning and suppression of breadth. Terminally exhausted CD8 T cells were present in broad neutralizers reinforcing the consequence of this antigenic load on the functional cellular immune response. Narrow neutralizers with lower viral load maintained progenitor exhausted CD8 T cells, identified as correlates of post-intervention viral control and long-term non-progression^63–69^. This implicates viral load-driven exhaustion trajectory as both an enabler of breadth and a critical determinant of functional immune responses to HIV infection and sensitivity to immune-based cure strategies. Together, these findings reframe neutralization breadth not as a product of effective immunity, but as a consequence of its failure.

## Results

### Cohort description

To profile the natural immune response to HIV infection and identify correlates of antibody breadth we profiled 27 treatment-naive PLWH for plasma neutralization breadth using samples collected around five years post-infection (Fig. 1A and B). For each individual, two longitudinal peripheral blood mononuclear cell (PBMC) samples, one collected on day 14 - 67 post-infection and the second around five years post-infection (matching plasma sample timing), were available for multi-omic profiling. After thawing and exclusion for low viability n = 16 early and n = 18 late samples passed quality control and were profiled by scRNA-seq and CyTOF; each CyTOF sample was profiled in both stimulated (viral-like TLR agonists) and unstimulated conditions (Fig. 1A-D, Fig. S1A). An additional six HIV-negative individuals from the same cohort were profiled by CyTOF and scRNA-seq.

**Fig. 1:**
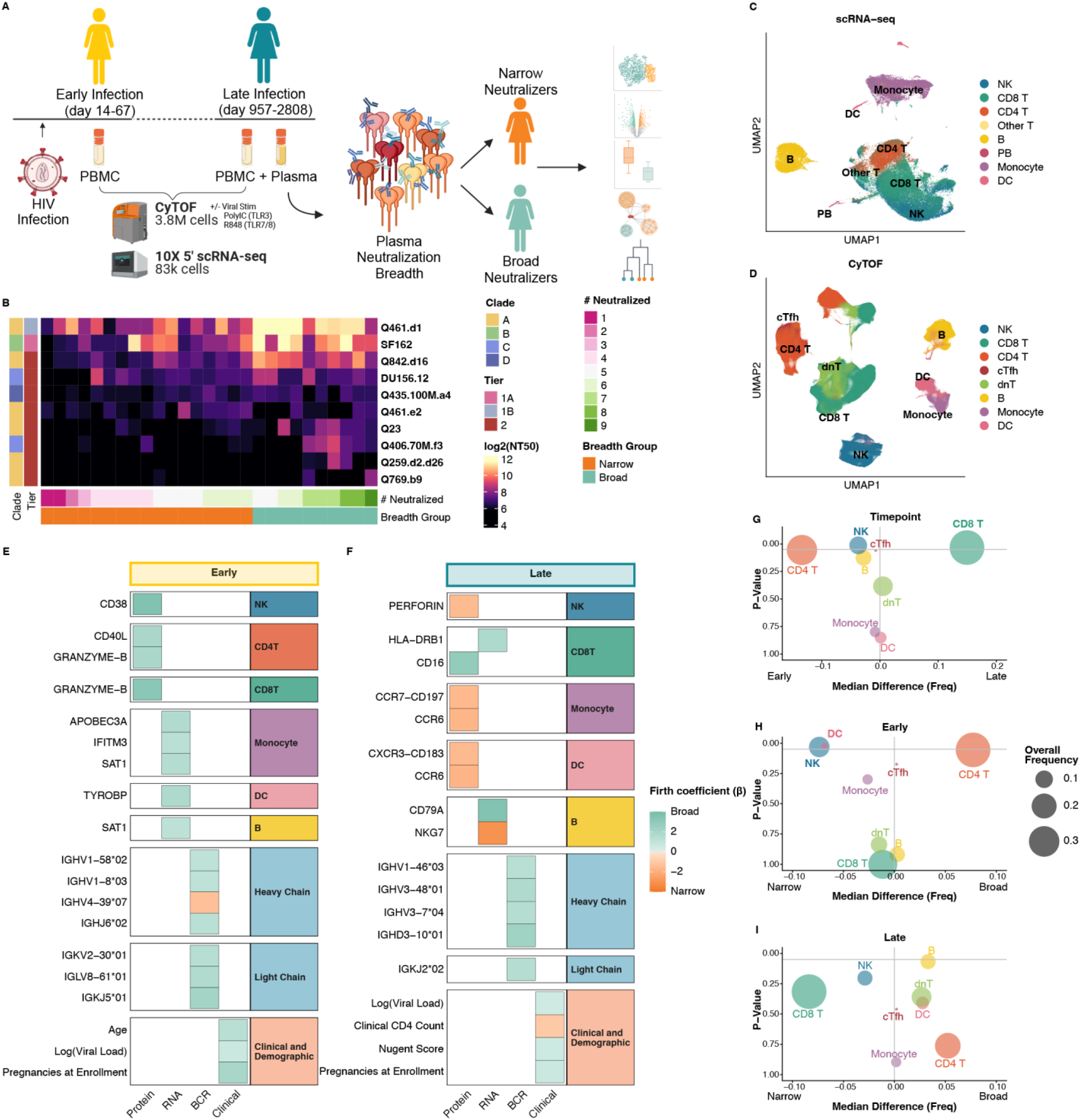
A comprehensive single-cell data set of untreated HIV infection with paired antibody breadth. **A.** Outline of samples and analysis plan. **B.** Heatmap of NT50 against 10 HIV pseudoviruses (row) for each subject (column). Each cell represents the average of 3 technical replicates. Columns are annotated by the number of pseudoviruses neutralized and breadth group. Row annotations indicate the tier and clade of each pseudovirus. **C.** UMAP of complete scRNA-seq data set colored by broad cell types. **D.** UMAP of complete CyTOF dataset colored by broad cell types. **E** and **F.** Heatmap of Stabl selected features in early (**E**) and late (**F**) infection from variable selection within each modality. Color represents z-score of post-hoc glm coefficient estimate (within modality) with positive values associated with narrow neutralizers and negative values with broad neutralizers. **G-I**. Volcano plot of donor-level median difference in cell frequencies (% of PBMC) in early and late infection (**G**), broad and narrow neutralizers in early infection (**H**), and broad and narrow neutralizers in late infection (**I**). P-values from wilcoxon rank sum test are displayed on the y-axis with significant differences bolded.

### Plasma neutralization of HIV viruses circulating in Kenya is variable

Antibody breadth was determined at the late infection timepoint by plasma neutralization assays in TZM-bl cells using a panel of 10 HIV pseudoviruses with patient-derived envelope sequences representing clades A, B, C, and D (Fig. 1B). The panel was majority clade A, reflecting the predominant circulating clade in Kenya to measure breadth against relevant strains^70,71^. The majority were tier 2 viruses, with tier 1A and 1B viruses included to capture the full range of neutralization sensitivity^72^. Together, this panel encompasses substantial antigenic diversity as the clades differ by approximately 30% in amino acid sequence^73,74^. Individuals neutralized 1 to 9 pseudoviruses, showing large variation in neutralization responses in this cohort. As expected, the majority of individuals neutralized tier 1A and B strains with tier 2 strains showing more variable susceptibility. To classify individuals, we performed principal component analysis (PCA) on NT50 values followed by k-means clustering (k = 2) to separate individuals into broad and narrow neutralizers (Fig. S1B). Narrow neutralizers neutralized 1 to 6 viruses and broad neutralizers neutralized 5 to 9 viruses with greater tier 2 neutralization. These groups allowed us to identify single cell transcriptomic (Fig. 1C) and proteomic (Fig. 1D) predictors of broad and narrow antibody responses.

### NK and CD4 T cells are re-modeled by HIV infection

To prioritize our analyses, we used Stabl, a supervised machine learning framework designed for high dimensional data where the number of features exceeds the number of observations^75^. Stabl combines subsampled elastic net regularized regression with artificial noise injection to estimate the frequency at which each experimental variable is selected above noise variables. For each modality (protein by CyTOF, RNA by scRNA-seq, BCR features from 5’ BCR amplification, and clinical records) and timepoint separate models were trained (Fig. 1E and F, Table S4 and S5). To assess direction and magnitude of association, post-hoc univariate Firth penalized logistic regression was performed for each selected feature; positive coefficients indicate enrichment in narrow neutralizers (orange) and negative coefficients indicate enrichment in broad neutralizers (teal). In early infection, markers of activation were associated with broad neutralization (which was measured at the late time point as discussed above) (Fig. 1E). In the protein model, NK cell expression of CD38 as well as CD8 T cell expression of granzyme-B were associated with broad neutralizers. In CD4 T cells, CD40L, which mediates B cell help, was associated with broad neutralizers in addition to granzyme-B expression, consistent with expansion of CD4+ cytotoxic T lymphocytes (CD4 CTL) in HIV infection^76^. Most selected RNA features were derived from monocytes with inflammatory gene *SAT1* and antiviral ISGs *IFITM3* and *APOBEC3A* (also a well characterized HIV restriction factor)^77^ all associated with broad neutralizers. Several heavy and light chain genes were associated with broad neutralizers including *IGHJ6*02* used in the PC39 bnAb and *IGHV1-8*03* which is associated with V2-apex bnAbs^78,79^. Viral load was also associated with broad neutralizers suggesting that antigenic pressure is associated with the observed widespread immune activation and breadth. Other clinical variables selected (number of previous pregnancies and age) were all associated with broad neutralizers.

In late infection, CD8 T and B cell activation remained associated with broad neutralizers (Fig. 1F). CD8 T cell protein expression of CD16, which mediates antibody-dependent cellular cytotoxicity (ADCC), and RNA expression of *HLA-DRB1,* indicating activation, were associated with broad neutralizers. *CD79A*, indicative of BCR signaling, in B cells was also associated with broad neutralizers. Heavy chain allele *IGHV1-46*03* was associated with broad neutralizers; the IGHV1-46 alleles are widely used in bnAbs^80–82^. Additionally, *IGKV1-33*01* light chain allele was associated with broad neutralizers and is used in some VRC01 class antibodies^83^. In narrow neutralizers, NK cytotoxicity indicated by perforin expression was selected in late infection, highlighting potential for Tfh cell pruning. CCR6, CCR7, and CXCR3 expression in monocytes and DCs, which are drivers of migration and inflammation, were additionally associated with narrow neutralizers. In B cells, *NKG7* was associated with narrow neutralizers which is not a typical B cell related gene but was reported to be expressed in B cells during chronic SIV infection^84^ and was expressed in cells with a BCR sequence in this dataset (Fig. S1C). Higher clinical CD4 T cell count was also associated with narrow neutralizers indicating a relative maintenance of CD4 T cell levels in this group, while viral load remained associated with broad neutralizers along with age and nugent score (bacterial vaginosis).

There were differences in cell type proportions across infection timepoints, with CD4 T cells and NK cells being more abundant in early infection and CD8 T cells in late infection (Fig. 1G). There were also timepoint-specific differences between broad and narrow neutralizers. In early infection, NK cells were significantly more abundant in broad neutralizers, while CD4 T cells were elevated in the narrow group, both consistent with variable selection (Fig. 1H). In late infection, there was an increased proportion of CD8 T cells in broad neutralizers that does not reach significance (Fig. 1I). Similar results were observed in more granular cell subsets (Fig. S1D-F).

### Broad neutralizers have increased viral load and lower CD4 Counts

To further understand associations between broad neutralization, pregnancies, age, higher plasma viral load, and lower CD4 T cell count, we examined clinical data across the full course of HIV infection in broad and narrow neutralizers. Broad neutralizers were older and had more previous pregnancies than narrow neutralizers, though no pregnancies occurred during the study period (Fig. S1G-I). At the late infection timepoint profiled, participants in the broad neutralizer group had significantly lower CD4 T cell count (Fig. 2A).

**Fig. 2:**
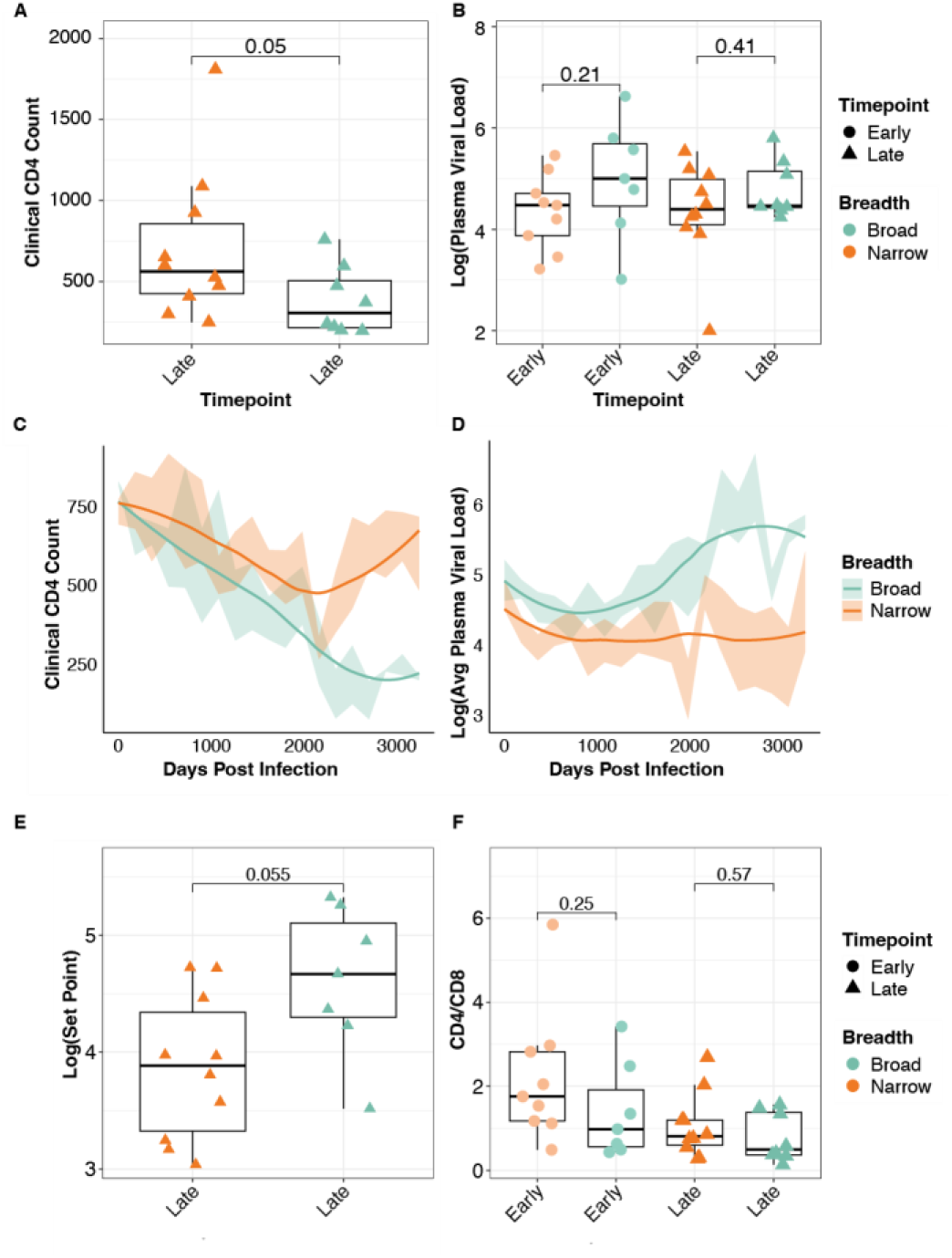
Broad neutralizers have lower CD4 count and higher viral load. **A.** Boxplot quantifying clinical CD4 T cell count in broad and narrow neutralizers at late infection timepoint. **B.** Boxplot quantifying log transformed viral load in broad and narrow neutralizers in early and late infection. **C** and **D.** Trajectory of log transformed viral load (**C**) and clinical CD4 count (**D**) in broad and narrow neutralizers in HIV infection. Data were grouped in 6-month intervals due to inconsistent visit timing. Shaded region represents 95% confidence interval; lines represent loess-smoothed trajectory. **E.** Boxplot quantifying log transformed viral set point broad and narrow neutralizers at late infection timepoint. **F.** Boxplot quantifying CD4 to CD8 ratio calculated using CyTOF data in broad and narrow neutralizers in early and late infection. P-values by two-sided Wilcoxon rank-sum test with Bonferroni’s correction for multiple hypothesis testing. Each point represents one donor.

In early infection, broad neutralizers had elevated viral loads, though this difference was not significant (Fig. 2B). However, when we examined the longitudinal clinical data for each individual, we found that broad neutralizers had a steeper decline in CD4 T cells and lower CD4 T cell count across infection (Fig. 2C). There was a corresponding higher viral load in broad neutralizers with an increase in viral load over time while narrow neutralizers stayed more constant (Fig. 2D). Broad neutralizers also had a trend towards higher viral set point and lower CD4 to CD8 ratio in early infection (Fig. 2E and F). To address whether demographic features, viral load, and CD4 T cell count confounded the observed immune associations with breadth, confounder-adjusted Firth penalized logistic regression was performed for each Stabl-selected feature. Adjusting for age, number of pregnancies, CD4 count and viral load did not attenuate the majority of selected associations, and many strengthened after adjustment, indicating negative confounding (Fig. S2). Effect size estimates were further robust to donor bootstrap resampling (Fig. S3). These data are consistent with previous reports that broad neutralizers have more severe HIV infection with higher viral load and greater CD4 T cell decline, and demonstrate that the identified immune associations with breadth are not purely attributable to demographic confounders, together validating this cohort as an appropriate setting to leverage single-cell, multi-omic data to explore underlying mechanisms and associations.

### NK cell activation and function diverge between broad and narrow neutralizers

In early infection, NK cells from broad neutralizers were characterized by markers of activation and proliferation. CyTOF profiling revealed significantly higher expression of CD38, HLA-DR, and Ki-67 and elevated expression of CD16 and granzyme-B compared to narrow neutralizers (Fig. 3A). Despite this activated state, broad neutralizer NK cells were functionally impaired. Stimulation of donor PBMCs with viral-like cocktail of TLR agonists revealed that upregulation of CD107a, CD8A, TRAIL, and TNFα was significantly greater in narrow neutralizer’s NK cells compared to broad neutralizers, indicating a higher functional response to viral-like signals in early infection (Fig. 3B). This discrepancy between activation and function in broad neutralizers prompted investigation of exhaustion; an aggregated transcriptomic exhaustion score (*PDCD1, HAVCR2, TIM3, TIGIT, EOMES*) confirmed that broad neutralizer NK cells had elevated exhaustion at the early timepoint, which became significant in late infection (Fig. 3C). In late infection, narrow neutralizers expressed significantly higher baseline levels of multiple cytotoxic proteins (Perforin, CD107A, TRAIL) exhibiting degranulation and the death receptor TRAIL (Fig. 3D). They also had significantly higher expression of CXCR5, the receptor that allows migration to lymph nodes^85–87^ where NK cells may kill Tfh cells, as well as CCR6. Broad neutralizers, by contrast, showed signs of progressive dysfunction not only in scRNA-seq exhaustion score, but also in accumulation of CD56 negative (CD56^neg^) NK cells, a hallmark of dysfunction in chronic infection (Fig. 3E).

**Fig. 3:**
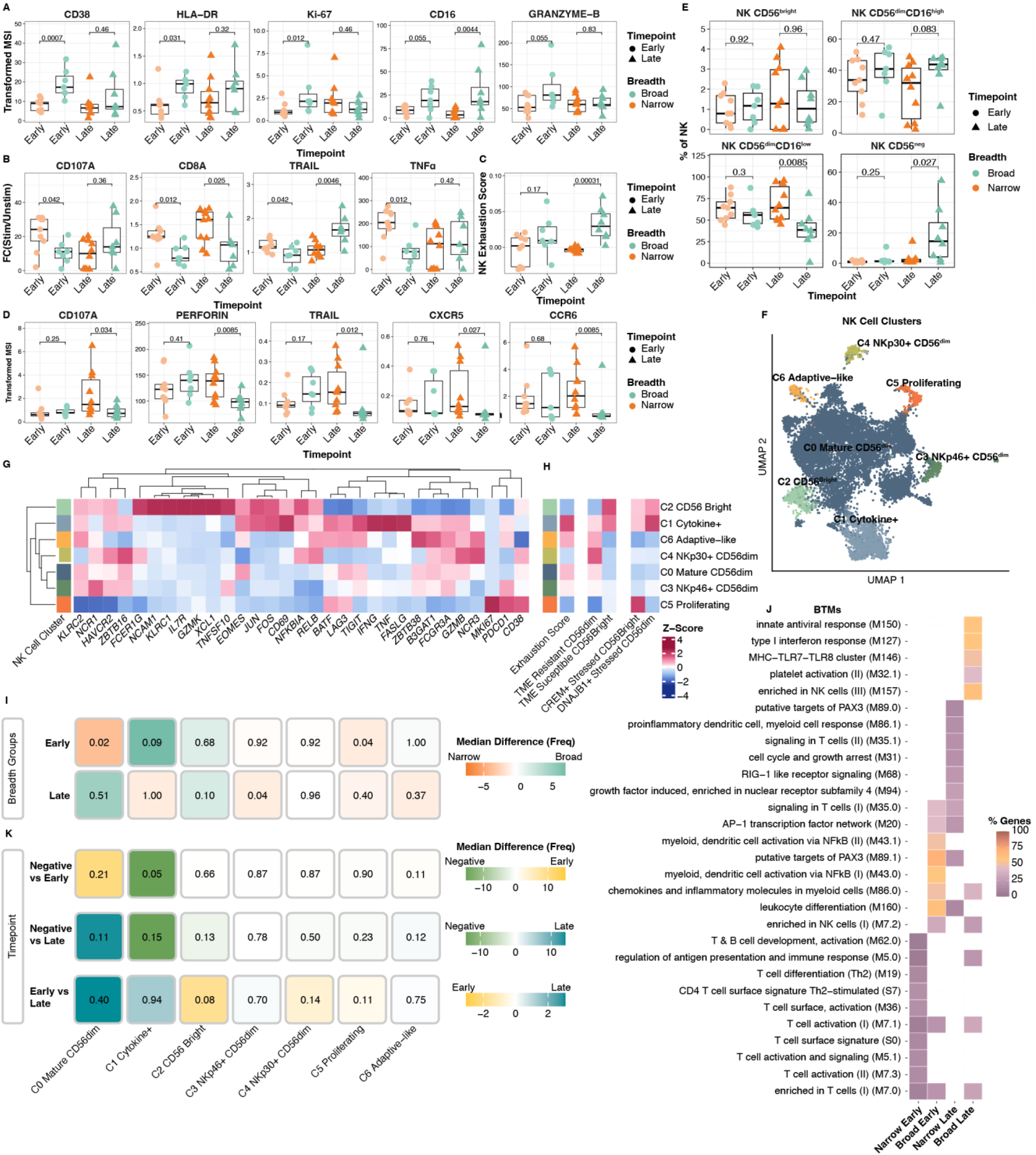
Early NK cell activation is followed by NK exhaustion in broad neutralizers. **A.** Boxplots quantifying arcsinh-transformed average expression in breadth groups of markers in NK cells from CyTOF dataset. **B.** Boxplots of fold-change (FC) in per-subject average expression in stimulated vs. unstimulated samples for each breadth group of markers in NK cells from CyTOF dataset. **C.** Boxplot of average NK exhaustion score (defined as expression of *LAG3*, *PDCD1*, *HAVCR2, TIGIT, EOMES*; see Methods) in breadth groups of NK cells from scRNA-seq dataset. **D.** Boxplots quantifying arcsinh-transformed average expression breadth groups of markers in NK cells from CyTOF dataset. **E.** Boxplots quantifying proportion of NK subsets manually gated (Fig. S9) in CyTOF dataset in breadth groups. **F.** UMAP of NK cells in scRNA-seq dataset colored by cluster annotation. **G.** Heatmap of normalized expression of NK cell genes in each annotated cluster ordered by hierarchical clustering. **H.** Heatmap of normalized expression of NK cell TME stress and exhaustion-related gene scores. **I.** Heatmap of median difference in frequency of each NK cell cluster in breadth groups, with associated p-values. **J.** Heatmap of top 10 BTM pathways enriched in each breadth group by over-representation analysis implemented with a one-sided hypergeometric test with FDR adjusted p < .05. Color indicates the proportion of module genes present in enriched gene lists. **K.** Heatmap of median difference in frequency of each NK cell cluster in infection timepoints, with associated p-values. P-values by two-sided Wilcoxon rank-sum test with Bonferroni’s correction for multiple hypothesis testing. Each point represents one donor.

To resolve the transcriptional correlates of these functional differences, we identified 7 NK cell clusters in scRNA-seq data (Fig. 3F and G). Transcriptional signatures of NK cell exhaustion and stress observed in tumor infiltrating NK cells (Tang et al. Cell. 2023., Netskar et al. Nat Immunol. 2024.) and exhaustion score described above was applied to each cluster to contextualize the dysfunction observed in broad neutralizers across other settings of chronic inflammation^88,89^. Finally, recently established NK1, NK2, and NK3 subset gene signatures (Rebuffet et al. Nat Immunol. 2024.) were also applied to facilitate cross-study comparison within a standardized framework of healthy NK cell heterogeneity ^90^ (Fig. S4A). C0 represented mature CD56^dim^ NK cells with expression of both *FCGR3A* (CD16) and *NCAM1* (CD56) with moderate expression of *B3GAT1* (CD57) and *KLRC2* (NKG2C) indicating maturity but lacking other markers of adaptive NK cells. C1 were cytokine-secreting NK cells with high expression of both *IFNG* and *TNF*, activation markers *CD38* and *CD69,* and the highest exhaustion score. C2 comprised CD56^bright^ NK cells with high expression of *NCAM1* along with markers of immaturity like *XCL1*, *IL7R*, and *GZMK* that shared features with CD56bright tumor susceptible NK cells. C3 and C4 represented CD56^dim^ NK cells differentiated by high expression of *NCR1* (NKp46) or *NCR3* (NKp30) respectively. C5 were proliferating NK cells with high expression of *MKI67* (Ki-67). C6 constituted adaptive-like NK cells with high expression of *B3GAT1, KLRC2,* and *ZBTB38* and low expression of *ZBTB16* and *FCER1G;* this cluster also had the highest expression of NK3 adaptive-like genes^91^.

Cluster composition in each group mirrored their functional phenotypes from protein data. In early infection, broad neutralizers had a trend for an elevated proportion of C1 activated, cytokine secreting NK cells (Fig. 3H, Fig. S4B). Consistent with premature activation and dysfunction observed in protein data, C1 scored highly in exhaustion as well as signatures of NK stress in tumors. Narrow neutralizers had a significantly greater proportion of C0 mature CD56^dim^ and C5 proliferating NK cells, both clusters were less exhausted, aligning with the increased function observed in narrow neutralizers in protein data. Using differentially expressed genes (DEGs) from NK cells in early or late infection, we performed gene set enrichment analysis using blood transcription modules (BTMs) described in Shuzhao et al. 2014., identifying modules significantly overrepresented in each group^92^ (Fig. 3I). In early infection, narrow neutralizers were enriched for general lymphocyte activation pathways, mirroring the mature NK cell phenotypes of C0 and C5. Broad neutralizers, by contrast, were enriched for AP-1 and NF-κB signaling, suggesting a molecular basis for the activated and exhausted C1 phenotype which also exhibits NF-κB (*NFKBIA, RELB*) and AP-1 (*FOS, JUN, BATF*) signaling.

In late infection, broad neutralizers had a trend towards a greater proportion of C2 CD56^Bright^ NK cells which share features with the tumor microenvironment (TME) susceptible CD56^Bright^ NK cell phenotype identified in Netskar et al. This indicates that broad neutralizers are enriched for an immature NK cell state with distinct exhaustive features aligned with immunosuppression in the TME. Narrow neutralizers had a significantly greater proportion of C3 NKp46+ CD56^dim^ NK cells in late infection which also express *TNFSF10* (TRAIL) consistent with the cytotoxic phenotype observed in narrow neutralizers by CyTOF in late infection. BTM analysis showed that narrow neutralizers remained enriched for pathways of lymphocyte activation and AP-1/RIG-I like signaling in late infection. Broad neutralizers showed a shift toward innate inflammatory pathways in late infection with enrichment of innate antiviral response and type I interferon response modules. Together, these data indicate that NK cells in broad neutralizers are characterized by a more inflammatory, innate-skewed transcriptional program relative to the lymphocyte activation signature predominating in narrow neutralizers, especially in late infection. This was further reinforced with NK expression of CD38 in early infection as a top variable associated with broad neutralizers and perforin in late infection associated with narrow neutralizers, underscoring that NK cell signals dominate phenotypic differences between breadth groups.

Finally, we also examined NK cell dynamics across HIV-negative, early, or late timepoints to characterize where breadth groups diverge from overall infection trajectories (Fig. 3J, Fig. S4B). HIV negative individuals had a greater proportion of C1 cytokine secreting NK cells compared to early or late HIV infection, though this was only significant in early infection. C2 CD56^bright^ and C5 proliferating NK cells were decreased in late infection and C6 adaptive-like NK cells increased, representing maturation, though none of these trends reached significance. Taken together, these findings reveal a temporal divergence in NK cells between broad and narrow neutralizers: broad neutralizers mount an early activated but exhaustion-prone response, whereas narrow neutralizers sustain a functionally competent NK cell program that retains cytotoxic potential over the course of infection.

### Broad neutralizers exhibit greater CD4 T cell help sustained in late infection

Among CD4 T cells we profiled bulk CD4 T cells, circulating Tfh (cTfh) cells, and CD4 CTLs in CyTOF data, all of which are known to be altered by HIV infection. We did not find differences in the frequency of circulating Tfh cells (CD4+ CXCR5+), memory, or helper subtypes between broad and narrow neutralizers (Fig. 4A, Fig. S5A and B). Both conventional CD4T and cTfh cells exhibited differences in the proportion of CD4 CTLs (CD4+ Perforin+ Granzyme-B+) between breadth groups with elevated CD4 CTLs in broad neutralizers in early infection and narrow neutralizers in late infection (Fig. 4B and C, Fig. S5C and D). In early infection, CD4 T cells from broad neutralizers exhibited features of both activation and B cell help, with significantly higher expression of CD40L, HLA-DR, ICOSL and TNFα (Fig. 4D). In contrast, narrow neutralizers had greater expression of CD45RA indicating relative maintenance of naive cells, as well as significantly greater upregulation of TNFα and a trend towards greater upregulation of PD-1 in response to viral-like stimulation at this timepoint, reflecting robust responses compared to broad neutralizers (Fig. 4D and E). In late infection, broad neutralizers exhibited sustained activation with significantly higher CD38 expression. Although broad neutralizers had reduced cytokine responsiveness, they retained a trend toward greater upregulation of ICOSL, a critical driver of antibody breadth, suggesting maintenance of B cell help despite diminished anti-viral function.

**Fig. 4:**
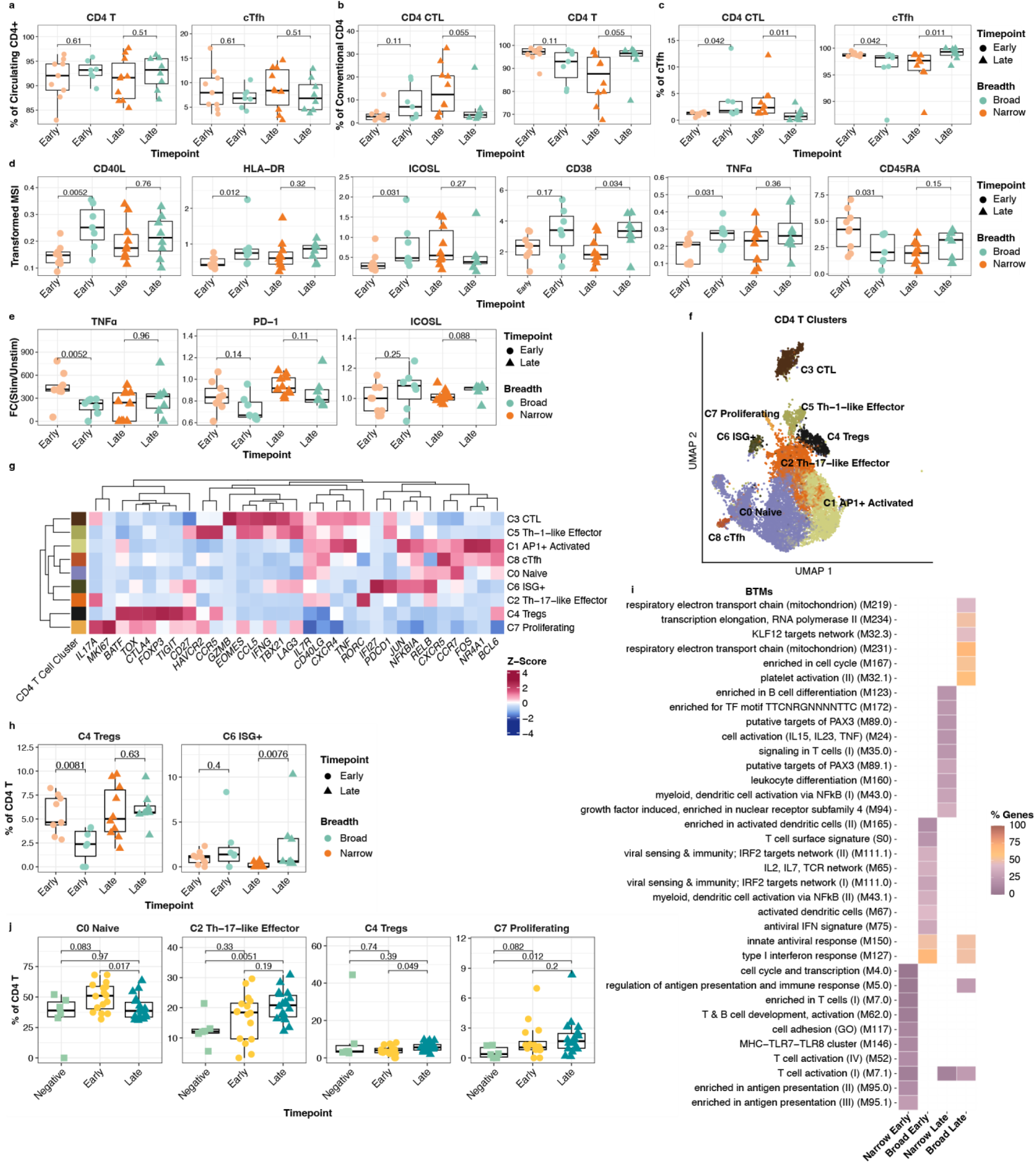
Broad neutralizers sustain CD4T cell help in late infection. **A.** Boxplots quantifying proportion of conventional CD4 T and circulating Tfh cells amongst all CD4 T cells from CyTOF dataset. **B.** Boxplots quantifying proportion of CD4 CTL and non-CTL amongst conventional CD4 T from CyTOF dataset. **C.** Boxplots quantifying proportion of CD4 CTL and non-CTL amongst circulating Tfh cells from CyTOF dataset. **D.** Boxplot of arcsinh transformed average expression in breadth groups of markers in CD4 T cells from CyTOF dataset. **E.** Boxplots fold-change (FC) in per-subject average expression in breadth groups of markers in CD4 T cells from CyTOF dataset. P-values by two-sided Wilcoxon rank-sum test with Bonferroni’s correction for multiple hypothesis testing. Each point represents one donor. **F.** UMAP of CD4 T cells in scRNA-seq dataset colored by cluster annotation. **G.** Heatmap of normalized expression of CD4 T cell genes in each annotated cluster ordered by hierarchical clustering. **H.** Heatmap of median difference in frequency of each CD4 T cell cluster in breadth groups, with associated p-values. **I.** Heatmap of top 10 BTM pathways enriched in each breadth group by over-representation analysis implemented with a one-sided hypergeometric test with FDR adjusted p <.05. Color indicates proportion of module genes present in enriched gene list. **J.** Heatmap of median difference in frequency of each CD4 T cell cluster in infection timepoints, with associated p-values. P-values by two-sided Wilcoxon rank-sum test with Bonferroni’s correction for multiple hypothesis testing. Each point represents one donor.

We identified 9 transcriptomic clusters among CD4 T cells (Fig. 4F and G). C0 represented naive CD4 T cells with high expression of *CCR7* and *IL7R*. C1 comprised activated CD4 T cells expressing AP-1 transcription factors, *CD69*, and *TNF*. C2 were Th17 CD4 T cells with expression of *RORC* and *IL17A*. C3 contained CD4 CTLs indicated by expression of *GZMB*. C4 were Tregs with expression of *FOXP3*. C5 contained Th1 CD4 T cells with expression of *TBX21* and *IFNG*. C6 expressed ISGs. C7 were proliferating CD4 T cells with high expression of *MKI67.* Finally, C8 contained cTfh cells with expression of *CXCR5* as well as the transcription factor *BCL6.* In early infection, C4 Tregs were significantly more abundant in narrow neutralizers (Fig. 4H, Fig. S5E). Tregs are implicated in suppression of B cells in PLWH who do not make bnAbs providing a mechanistic link to narrow neutralizers^14^. No clusters were enriched in broad neutralizers in early infection, however, BTM analysis revealed enrichment of pathways of interferon response, potentially a consequence of higher viral loads in this group (Fig. 4I). Furthermore, in late infection, C8 ISG+ CD4 T cells were significantly more abundant in broad neutralizers aligned with BTM enrichment of anti-viral pathways and higher viral loads.

Infection associated changes in cluster composition included a significant decrease in C0 naive CD4 T cells in late infection, compared to early infection indicating maturation (Fig. 4J, Fig. S5F). This was reinforced by a significantly greater proportion of C2 Th17-like and C7 proliferating CD4 T cells in late infection compared to HIV-negative individuals. Tregs were also significantly increased in late infection, compared to early infection, indicating that the overall trajectory of HIV disease is distinct from that in narrow neutralizers that had an early accumulation of Tregs. Taken together, these data reveal that broad neutralizers are characterized by a more activated, helper-competent CD4 T cell response present in early infection and sustained into chronic disease, with elevated ICOSL suggesting preserved B cell help capacity. In contrast, narrow neutralizers show early preservation of a naive phenotype, greater anti-viral cytokine secretion, and early Treg accumulation.

### Broad neutralizers display early B cell activation that wanes over the course of HIV infection

In early infection, B cells from broad neutralizers display greater activation with significantly higher protein expression levels of PD-1 and TNFα, as well as elevated levels of IFNγ (Fig. 5A). Broad neutralizers also had significantly lower proportions of naive B cells and higher expression of IgM, but not IgD, which could indicate early stages of class-switching (Fig. 5A and B). To further characterize B cell transcriptomic responses, we identified 8 clusters of B cells in transcriptomic space (Fig. 5C). All clusters contained B cells with successfully sequenced BCR and varying levels of somatic hypermutation (Fig. 5D and S6A). Naive B cells were represented in C0, identified by their expression of *IGHM*, *IGHD*, *IL4R*, *TCL1A*. C1 were atypical B cells with expression of *TBX21* (T-bet), *LILRB1*, *IL10*, and *ITGAX* (CD11c); these are tissue-like memory cells that expand during chronic infection. C2 consisted of memory B cells with expression of *CD27*. C3 were activated B cells with expression of AP-1 transcription factors as well as *TNF*. C4 comprised mature B cells with expression of *AIM2*, *BANK1*, and *HLA-DR*. C5 were plasmablasts (PBs) with expression of *XBP1*, *MZB1, MKI67* and high *CD27.* C6 were exhausted atypical B cells that expressed exhaustion markers *LAG3* and *TIGIT* as well as activation markers *CCL5,* and *IFNG.* C7 contained atypical B cells with high expression of *NFKBIA* (the gene that encodes NF-κB) (Fig. 5E).

**Fig. 5:**
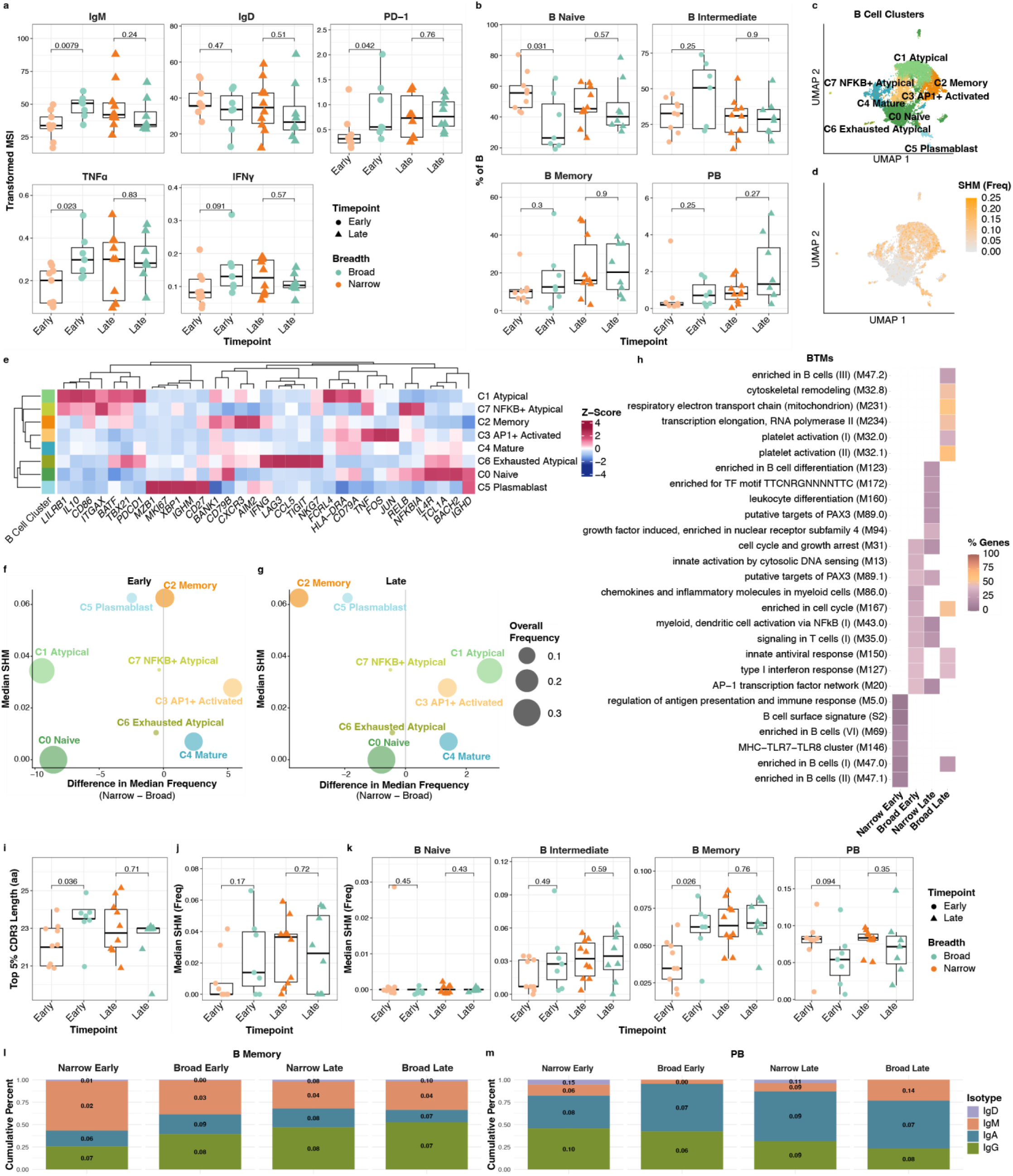
Broad neutralizers exhibit early but sub-optimal B cell responses. **A.** Boxplot of arcsinh transformed average expression in breadth groups of markers in B cells from CyTOF dataset. **B.** Boxplots quantifying proportion of B cell subsets in breadth groups from CyTOF dataset. **C.** UMAP of B cells in scRNA-seq dataset colored by cluster annotation. **D.** UMAP of B cells in scRNA-seq dataset colored by somatic hypermutation (SHM) frequency. **E.** Heatmap of normalized expression of B cell genes in each annotated cluster ordered by hierarchical clustering. **F** and **G**. Volcano plot of median difference in cluster frequencies (% of B) between broad and narrow neutralizers in early (**F**) and late (**G**) infection. Y-axis shows the median SHM of each cluster. **H.** Heatmap of top 10 BTM pathways enriched in each breadth group by over-representation analysis implemented with a one-sided hypergeometric test with FDR adjusted p < .05. Color indicates the proportion of module genes present in enriched gene lists. **I.** Boxplot of per-donor median length of top 5% of CDR3 sequences in breadth groups. **J.** Boxplot of per-donor median SHM in breadth groups. **K.** Boxplots of per-donor median SHM from each B cell subset in breadth groups. P-values by two-sided Wilcoxon rank-sum test with Bonferroni’s correction for multiple hypothesis testing. Each point represents one donor. **L** and **M.** Cumulative bar plots of immunoglobulin heavy chain gene usage from memory B cells (**L**) and plasmablasts (**M**) in breadth groups colored by isotype with median SHM of each isotype displayed.

Cluster abundances in breadth groups revealed distinct early and late infection phenotypes by breadth group (Fig. 5F and G, Fig. S6B). In early infection in broad neutralizers, C3 AP-1+ B cells were elevated, though this difference was not significant. This cluster expressed moderate levels of *CD86* and *HLA-DRA* indicating functional B cell responses as well as AP-1 and NF-κB signaling genes, consistent with pathway analysis which demonstrated enrichment for BTMs related to AP-1 signaling, NF-κB signaling, and interferon response (Fig. 5H). In narrow neutralizers, C6 exhausted atypical B cells were significantly more abundant in early infection. This cluster expressed markers of both activation and exhaustion as well as *NKG7*, the greatest predictors of narrow neutralization in variable selection. *NKG7*, in particular, suggests that this is an aberrant, dysfunctional cell type associated with chronic infection. C5 PBs were also elevated indicating that this dysregulated response is accompanied by premature differentiation, consistent with BTM enrichment for B cell activation and antigen presentation in early infection in narrow neutralizers. In late infection, C2 memory B cells and C7 NF-κB+ atypical B cells were elevated in narrow neutralizers. These C2 memory B cells expressed T-bet but not other markers of atypical B cells suggesting a transition towards atypical, dysfunctional cell state. No clusters were elevated in late infection in broad neutralizers; most cells are C1 atypical B cells, indicative of widespread dysfunction at this timepoint.

BCR features (somatic hypermutation (SHM), CDR3 length etc.) were also derived from B cells using 10X 5’ capture. BCR analysis further supported the divergence between breadth groups in early infection. In early infection broad neutralizers have higher SHM as well as longer CDR3 sequences in total B cells; however, this effect was not observed in late infection at the time when breadth was measured (Fig. 5I and J). This difference in SHM in early infection was driven specifically by memory B cells that had significantly higher SHM in broad neutralizers in early infection; PBs from broad neutralizers paradoxically showed lower SHM than narrow neutralizers, indicating inadequate responses in cell subsets directly responsible for antibody secretion (Fig. 5K). This aligned with high SHM in clusters associated with narrow neutralizers in late infection. Isotype usage revealed that memory B cells from broad neutralizers in early infection had greater IgG usage and greater average SHM in IgA, reflecting early increased class switching and maturation in the memory compartment (Fig. 5L). In PBs, broad neutralizers had greater IgA usage in early infection while narrow neutralizers had greater IgG use as well as greater SHM in IgG with other B cell subsets having fewer differences between broad and narrow neutralizers (Fig. 5M, Fig. S6D and E). Together, these BCR-level findings point to early infection as a critical window distinguishing neutralization phenotype. Broad neutralizers exhibit a more robust early memory B cell response, but this difference is diminished by late infection as the overall B cell response wanes and atypical B cell phenotype dominates.

This was also observed in the overall shift in B cell phenotypes in HIV infection overall. AP-1+ B cells were elevated in HIV-negative individuals indicating a decrease in functional B cells during infection which is distinct from the elevated AP-1+ B cells in broad neutralizers in early infection (Fig. S6C). C1 atypical B cells were significantly more abundant in late infection compared to HIV-negative individuals and elevated compared to early infection; C2 memory B cells were also elevated over the course of infection indicating accumulation of mature B cells. C6 exhausted atypical and C7 NF-κB+ atypical B cells were narrow neutralizer specific and were not significantly changed over the course of infection. Collectively, these data suggest that the B cell compartment undergoes substantial functional decline over the course of HIV infection, with the early response representing a key determinant of long-term neutralization breadth.

### Terminally exhausted CD8 T cells are enriched in broad neutralizers

The activation observed in broad neutralizers in early infection extended to CD8 T cells, with significantly greater protein expression of CD161, Granzyme B, and TNFα indicating cytolytic activity and inflammation (Fig. 6A). This activation was accompanied by maturation, reflected by significantly lower CD45RA expression and a shift in CD8 T cell subset frequencies. CD8 T Effector Memory (EM) were significantly elevated and CD8 Naive depleted in broad neutralizers in early infection (Fig. 6B). CD8 T cells in broad neutralizers also expressed greater levels of ICOSL in early infection and ICOS in late infection. These costimulatory proteins are not typically associated with CD8 T cells but are reported to be upregulated in settings of chronic inflammation like cancer or HIV infection^93,94^. Despite their activated state, CD8 TEM from broad neutralizers had reduced functional responses upon stimulation with significantly lower upregulation of TNFα, CD107A, and ICOS as well as a trend towards lower PD-1 upregulation compared to narrow neutralizers indicating exhaustion in this specific cell subset. This impairment was restricted to CD8 TEM and was not observed in non-EM CD8 T cell subsets (Fig. 6C and D).

**Fig. 6:**
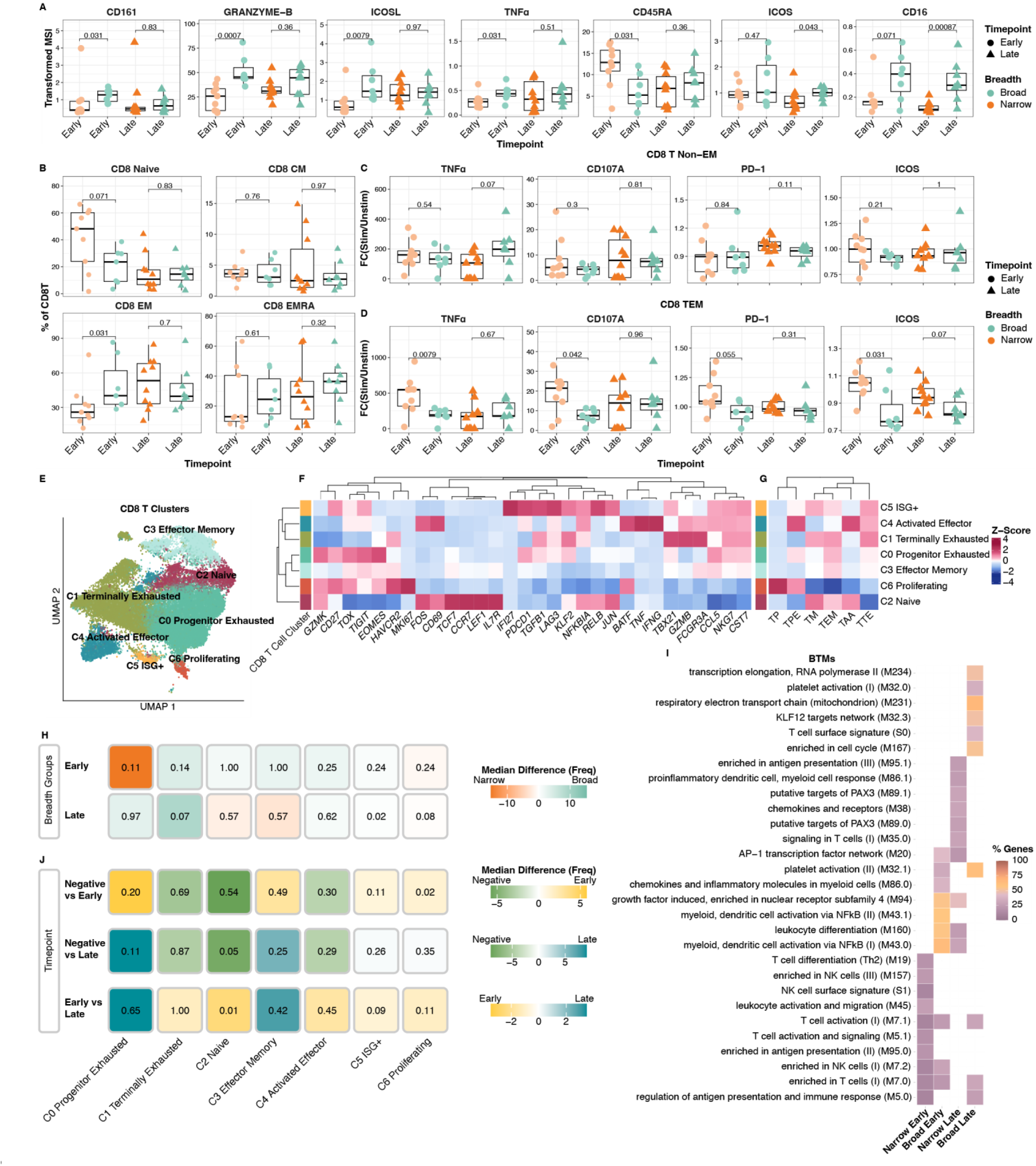
Broad neutralizers are enriched for terminally exhausted CD8 T cells. **A.** Boxplots quantifying arcsinh transformed average expression in breadth groups of markers in CD8 T cells from CyTOF dataset. **B.** Boxplots quantifying proportion of CD8 T cell subsets manually gated (Fig. S8) in CyTOF dataset in breadth groups. **C** and **D.** Boxplots fold-change (FC) in per-subject average expression in breadth groups of markers in non-effector memory (EM) (**C**) and EM (**D**) CD8 T cells from CyTOF dataset. **E.** UMAP of CD8 T cells in scRNA-seq dataset colored by cluster annotation. **F.** Heatmap of normalized expression of CD8 T cell genes in each annotated cluster ordered by hierarchical clustering. **G.** Heatmap of normalized expression of CD8 T cell exhaustion and activation module scores from Oliveira et al. Nature 2021. (TP = CD8 T proliferating, TPE = CD8 T progenitor exhausted, TM = CD8 T memory, TEM = CD8 T effector memory, TAA = CD8 T acutely activated, TTE = CD8 T terminally exhausted.) **H.** Heatmap of median difference in frequency of each CD8 T cell cluster in breadth groups, with associated p-values. **I.** Heatmap of top 10 BTM pathways enriched in each breadth group by over-representation analysis implemented with a one-sided hypergeometric test with FDR adjusted p < .05. Color indicates the proportion of module genes present in enriched gene lists. **J.** Heatmap of median difference in frequency of each CD8 T cell cluster in infection timepoints, with associated p-values. P-values by two-sided Wilcoxon rank-sum test with Bonferroni’s correction for multiple hypothesis testing. Each point represents one donor.

Chronic immune activation and CD8 T cell exhaustion observed in HIV infection shares features with tumor-infiltrating CD8 T cell biology where the transcriptional heterogeneity of exhaustion and function under chronic stimulation has been extensively characterized. Applying canonical CD8 T cell subset genes along with published gene signatures of terminally exhausted (TTE) and progenitor exhausted (TPE) gene signatures to our scRNA-seq data^95^, we identified 7 CD8 T cell clusters (Fig. 6E-G). C0 comprised progenitor exhausted CD8 T cells that expressed stress markers *TOX* and *TIGIT*, but retained expression of *GZMK* and *TCF7,* and scored highly on TPE signature, indicating preserved functional potential. C1 were terminally exhausted characterized by *LAG3* and *TIM3* expression and the highest TTE score among clusters. C2 represented naive CD8 T cells with expression of *IL7R*, *LEF1*, *CCR7*, and *GZMK* which also expressed AP-1 and NF-κB pathway genes. C3 contained CD8 TEM with expression of *CCL5* and *CTS7*. C4 comprised activated effector CD8 T cells that had the highest acutely activated (TAA) score and expressed *IFNG*, *TNF* and *CD69* and AP-1 signaling genes. C5 contained exhausted ISG+ CD8 T cells that had high effector and exhausted scores. C6 constituted proliferating CD8 T cells marked by *MKI67* expression with high proliferating (TP) and TPE scores.

In early infection, narrow neutralizers had an elevated proportion of C0 progenitor exhausted CD8 T cells consistent with relative maintenance of CD8 T cell function observed in CyTOF data (Fig. 6H, Fig. S7A).

In contrast, broad neutralizers had elevated proportions of C1 terminally exhausted CD8 T cells suggesting the robust activation state observed in protein data leads to early progression towards terminal exhaustion in broad neutralizers. This divergence in CD8 T cell composition and function was reflected in pathway analysis using BTMs, where broad neutralizers had enrichment of NF-κB and AP-1 pathways, while narrow neutralizers were enriched for general lymphocyte activation (Fig. 6I). This suggests that while both broad and narrow neutralizers are under chronic inflammatory pressure in early infection, diverging transcriptional programs are already shaping distinct functional trajectories at this early timepoint.

In late infection, narrow neutralizers showed elevated proportions of C6 proliferating CD8 T cells with high TPE scores and maintained enrichment of lymphocyte activation pathways, indicating preserved T cell function. In contrast, broad neutralizers had a significantly greater proportion of C5 ISG+ CD8 T cells with TTE phenotype and maintained elevated C1 terminally exhausted CD8 T, suggesting progressive enrichment of exhausted populations over the course of infection in broad neutralizers. C1 terminally exhausted CD8 T were highest expressors of *FCGR3A* (CD16) which mediates ADCC. Broad neutralizers expressed higher protein levels of CD16 in both early and late infection linking CD8 T cell exhaustion and ADCC function. Broad neutralizers in late infection switched toward enrichment of metabolic pathways, reflecting metabolic reprogramming associated with chronic activation and exhaustion.

Infection driven remodeling and effector cell maturation of the CD8 T cell compartment was evident over the course of HIV infection (Fig. 6J, Fig. S7B). C2 naive CD8 T cells significantly decreased over both early and late infection compared to HIV-negative controls. C5 ISG+ and C6 proliferating CD8 T cell clusters peak in early infection, further indicating that exhaustion programs arise early in infection. These data reveal a divergence in the CD8 T cell response between broad and narrow neutralizers. Broad neutralizers with greater antigen burden leads to sustained activation that ultimately promotes terminal exhaustion and functional impairment. In contrast, the more preserved progenitor exhausted state in narrow neutralizers with lower viral load could be a consequence of greater long-term functional capacity.

### MDSC-like monocytes are associated with broad neutralizers

Monocytes were activated in broad neutralizers in both early and late infection indicated by elevated levels of CD11c and HLA-DR in early infection and CD161 in late infection (Fig. 7A). Narrow neutralizers expressed significantly greater levels of CCR6, CCR7, and CXCR3 in late infection indicating a migratory phenotype also reflected in Stabl variable selection. In scRNAseq we found 8 clusters amongst monocytes with varying functional states (Fig. 7B). To further characterize the functional identity of each cluster, we scored all clusters for gene signatures associated with monocyte derived suppressor cells (MDSCs) and monocytes in sepsis^96,97^ (Fig. 7C and D). C0 were MDSC-like CD14 classical monocytes with high expression of *TGFB1* alongside other features of suppression *IL10*, *PDCD1*, and *ITGAM* (CD11B). C1 contained both intermediate (CD16+ CD14+) and CD16+ inflammatory monocytes with moderate expression of ISGs. C2 comprised cytokine-expressing CD14 monocytes (*TNF*, *IL1B*) as well as features of exhaustion and suppressive activity. C3 represented MDSC-like classical monocytes. C4 were activated CD14 monocytes characterized by high expression of *CD38* and *CD69*. C5 contained stressed CD14 monocytes expressing heat-shock proteins (HSPs). C6 were ISG+ inflammatory CD16 monocytes with moderate MDSC phenotype. C7 contained CD16 monocytes with complement expression (*C1QA*, *B*, *C*) with additional expression of ISGs.

**Fig. 7:**
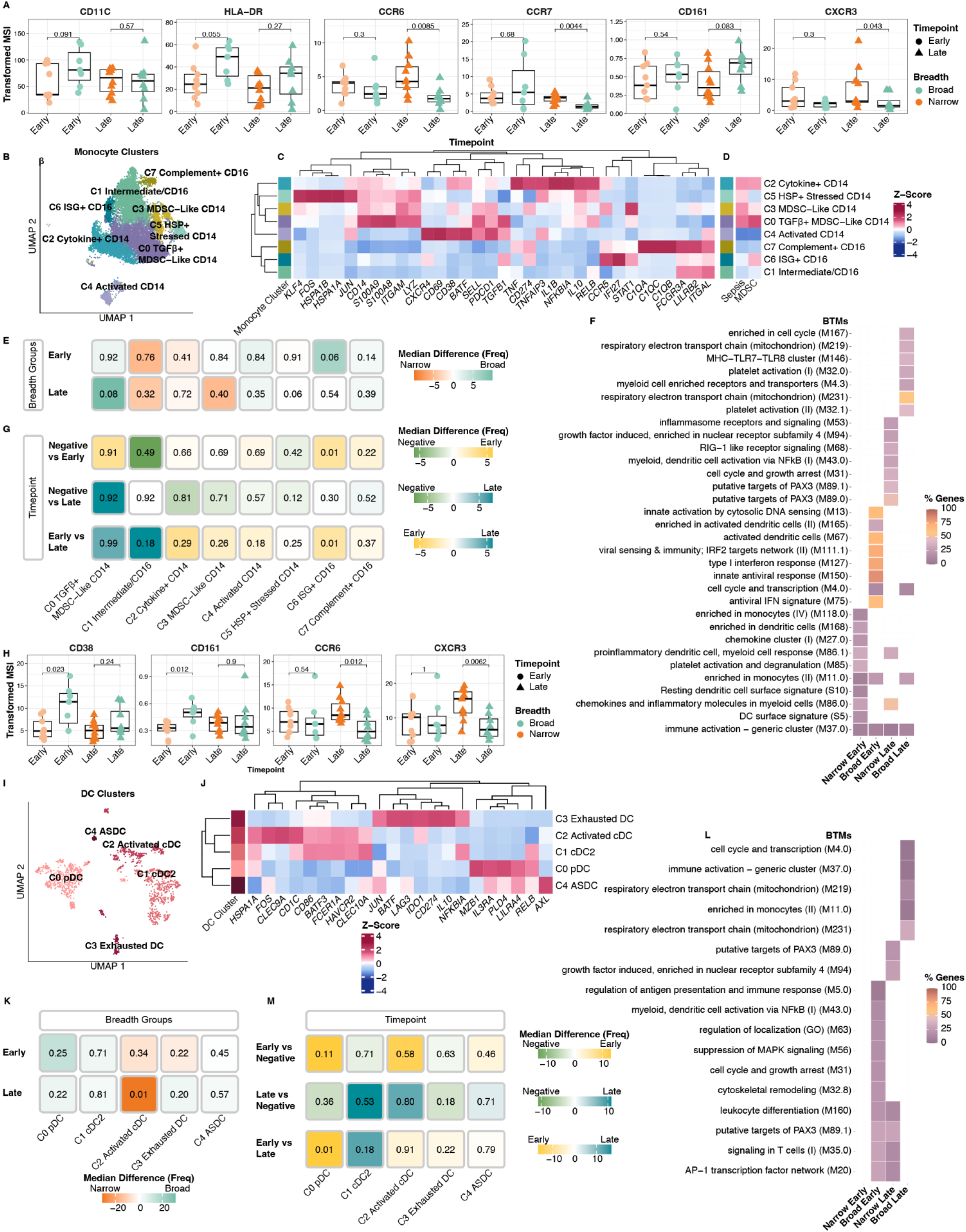
Myeloid compartment displays drivers of exhaustion and activation. **A.** Boxplot of arcsinh transformed average expression in breadth groups of markers in Monocytes from CyTOF dataset. **B.** UMAP of Monocytes in scRNA-seq dataset colored by cluster annotation. **C** and **D.** Heatmap of normalized expression monocyte genes (**C**) and monocyte MDSC and Sepsis score (**D**) in each annotated cluster ordered by hierarchical clustering. **E.** Heatmap of median difference in frequency of each Monocyte cluster in breadth groups, with associated p-values. **F.** Heatmap of top 10 BTM pathways enriched in each breadth group by over-representation analysis implemented with a one-sided hypergeometric test with FDR adjusted p < .05. Color indicates the proportion of module genes present in enriched gene lists. **G.** Heatmap of median difference in frequency of each Monocyte cluster in breadth groups, with associated p-values. **H.** Boxplot of arcsinh transformed average expression in breadth groups of markers in DCs from CyTOF dataset. **I.** UMAP of DCs in scRNA-seq dataset colored by cluster annotation. **J.** Heatmap of normalized expression DC genes in each annotated cluster ordered by hierarchical clustering. **K.** Heatmap of median difference in frequency of each DC cluster in breadth groups, with associated p-values. **L.** Heatmap of top 10 BTM pathways enriched in each breadth group by over-representation analysis implemented with a one-sided hypergeometric test with FDR adjusted p < .05. Color indicates the proportion of module genes present in enriched gene lists. **M.** Heatmap of median difference in frequency of each Monocyte cluster in infection timepoints, with associated p-values. P-values by two-sided Wilcoxon rank-sum test with Bonferroni’s correction for multiple hypothesis testing. Each point represents one donor.

In early infection, C6 ISG+ and C7 complement+ CD16 monocytes trended towards higher abundance in broad neutralizers (Fig. 7E, S8A). ISG expression in both clusters likely reflected higher viral loads driving type I interferon in broad neutralizers. This was corroborated by pathway analysis where monocytes from broad neutralizers were enriched for multiple interferon response pathways (Fig. 7F). While no clusters were elevated in narrow neutralizers in early infection, pathway analysis demonstrated enrichment for myeloid activation and migration, suggesting that the migratory potential observed by CyTOF in late infection may be initiated in early infection. In late infection, C0 MDSC-like monocytes, expressing *TGFB1*, *PDCD1*, and *IL10,* and sharing features with sepsis-mediated inflammation were elevated in broad neutralizers which could mediate the exhaustion observed in other immune compartments in late infection. C5 *HSP*+ stressed monocytes were elevated in narrow neutralizers in late infection. C5 expressed AP-1 transcription factors, low levels of *CD69*, *IL1B*, *TNF*, and *IL10*, and shared gene programs with monocytes in sepsis but not MDSCs indicating an inflammatory rather than suppressive phenotype in narrow neutralizers. Pathway analysis confirmed divergent phenotypes between monocytes in broad and narrow groups. Narrow neutralizers were enriched for activation pathways like RIG-1, NF-κB, and inflammasome indicating sustained innate immune activation whereas broad neutralizers were enriched for metabolic reprogramming pathways indicating the transition toward MDSC-like phenotype and functional suppression. HIV infection itself remodeled the monocyte compartment with C6 ISG+ CD16 Monocytes significantly more abundant in early infection, reflecting the initial antiviral response and immune activation (Fig. 7G, S8B). Together, these data demonstrate that monocyte phenotypes diverge between broad and narrow neutralizers, with broad neutralizers developing immunosuppressive MDSCs that may contribute to immune dysfunction, while narrow neutralizers maintain inflammatory monocyte populations capable of supporting adaptive immune responses.

### Broad neutralizers exhibit early dendritic cell activation followed by progressive exhaustion

Broad neutralizer dendritic cells (DCs) were more activated in early infection with significantly higher protein expression of CD38, and CD161 (Fig. 7H). In late infection, narrow neutralizers exhibited higher expression of CCR6 and CXCR3 indicating inflammation and migration. In scRNA-seq we identified 5 DC clusters (Fig. 7I and J). C0 were plasmacytoid DCs (pDCs) with expression of *IL3RA*, *PLD4*, and *MZB1*. C1 contained cDC2s with expression of *FCER1A*, *CLEC9A*, *CD1C*. C2 represented activated cDCs with expression of genes representative of both cDC1s and cDC2s. C3 were exhausted with expression of *IDO1*, *CD2749* (PD-L1), *IL10* and *LAG3*. C4 contained *AXL*+ DCs (ASDC).

In early infection, there were no significant differences in cluster composition, but C0 pDCs were slightly elevated in broad neutralizers which could be a source of interferon, aligning with ISG expression across other cell types at this timepoint (Fig. 7K, S8C). Consistent with broad DC activation, pathway analysis revealed enrichment of AP-1 signaling, NF-κB activation, and antigen presentation in broad neutralizers in early infection (Fig. 7L). In late infection, C0 pDCs remain elevated in addition to C3 exhausted DCs in broad neutralizers. Enriched pathways in this group indicate a metabolic shift suggesting a transition towards exhaustion. In contrast, activated cDCs were significantly more abundant in narrow neutralizers in late infection indicating maintenance of immune activation and innate-adaptive coordination supported by enriched pathways like AP-1 signaling. DC subsets also changed over the course of HIV infection. C0 pDCs peaked in early infection with a significant decrease in late infection, consistent with interferon signaling across the immune system in early infection (Fig. 7M, S8D). Broad neutralizers showed early DC activation followed by progressive exhaustion, whereas narrow neutralizers showed relative maintenance of DC-driven immune stimulation into late infection.

### Divergent cell-cell communication networks underlie distinct immune trajectories in broad and narrow neutralizers

To identify cell-cell communication pathways that differed between narrow and broad neutralizers, we leveraged the multinichenet package that predicts the top differentially expressed cell-cell communication pairs between groups based on ligand-receptor and downstream signaling gene expression, highlighting cell-cell communication pairs likely to be actively signaling from scRNA-seq data^98^. The top 25 cell-cell communication pairs are displayed and predicted interactions that are not mediated by extracellular proteins are colored transparent (Fig. 8A-C). In early infection, broad neutralizers had multiple predicted signaling pathways related to immunosuppression, consistent with robust immune activation followed by negative regulation (Fig. 8A). Multiple cell types were predicted to engage *LILRA4* on DCs via *BST2*. *BST2* is interferon stimulated, but signaling via LILRA4 dampens interferon secretion in pDCs, indicating that broad neutralizers mount a strong interferon response triggering inhibitory feedback^99,100^. *HLA-F,* which is also interferon induced, was predicted to engage inhibitory receptors *LILRB1/2*^101^ on monocytes and *KIR3DL2*^102^ on NK cells. LILRB1/2 engagement on monocytes promotes MDSC phenotype, consistent with monocyte phenotypes observed in broad neutralizers^103,104^. *HLA-DQB1* on B cells engaging *LAG3* on CD8 T cells further indicates suppression of cytotoxic responses ^105–107^. B cells were predicted to send signals related to pro-inflammatory IL-6 family cytokines; *EBI3* was predicted to engage *IL6ST* and *IL27RA* (shared receptors in the IL-6 family) in CD4 T cells and NK cells respectively^108^. Signaling via IL27RA can promote antiviral NK cell activity but also induces IL-10 and suppressive programs, consistent with the dual activation-exhaustion phenotype in early broad neutralizers^109,110^. In early infection, narrow neutralizers had fewer predicted interactions. Monocytes are predicted to send signals to DCs via *LGALS3*-*ANXA2*, an interaction associated with DC maturity, cross presentation, and NF-κB signaling^111^. Additional predicted interactions include *FN1* from monocytes engaging *CD44* on T cells which promotes Th1 polarization necessary for anti-viral immune responses^112,113^.

**Fig. 8:**
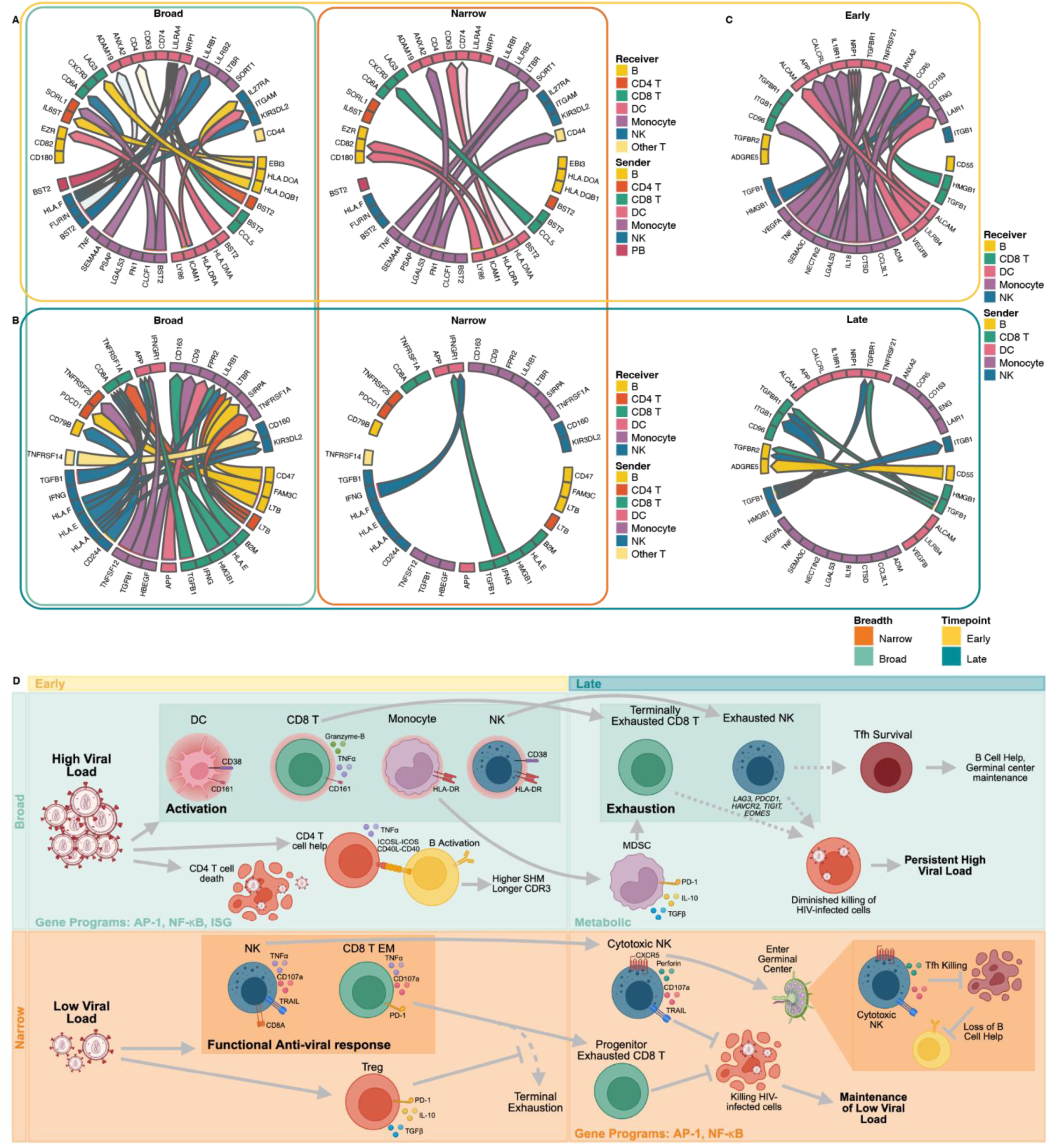
Cell-cell communication differs between breadth groups and infection timepoints. **A-C**. Circos plot showing top 25 validated predicted cell-cell communication pairs colored by sender cell split by broad and narrow neutralizers in early (**A**) and late (**B**) infection or timepoint (**C**). **D**. Proposed model of divergent immune responses in broad (top) and narrow (bottom) neutralizers in early (left) and late (right) HIV infection. Solid arrows indicate causal progression, dashed arrows indicate attenuated processes, and flat head arrows indicate inhibition, suppression or killing.

In late infection, broad neutralizers had predicted TNF signaling as well as inhibitory and immune checkpoint pathways (Fig. 8B). CD4 T cells were predicted to send signals via *LTB*, engaging *LTBR* and *TNFRSF1A* on monocytes and CD8 T cells respectively, with B cells similarly engaging LTBR on monocytes via LTB^114^. Reciprocally, monocytes were predicted to engage *TNFRSF25* on CD4 T cells via *TNFRSF12*^115^. Immune checkpoints were also predicted including *CD47* engaging *SIRPA* on monocytes and monocyte *LGALS3* engaging *LAG3* on NK cells driving suppression^116,117^. NK cells were predicted to drive activation in CD8 T cells via *HLA-F* and *HLA-A* and *HLA-E* engagement. Like in early infection, NK cells received inhibitory signals via KIR3DL2. In contrast, narrow neutralizers in late infection showed predicted *IFNG* secretion by CD8 T cells and NK cells, acting on DCs, consistent with DC maturation and cross presentation highlighting the sustained antiviral immunity in narrow neutralizers^118^.

To identify cell-cell communication patterns that shift with disease progression regardless of neutralization breadth, we compared predicted interactions enriched in early versus late infection (Fig.8C). Early infection was characterized by monocyte-centric inflammatory signaling. Monocytes were predicted to engage one another via *LGALS3*-*ENG* and *LGALS3*-*ANXA2*, reflecting autocrine monocyte activation and inflammatory amplification^111,119^. *CCL3L1*-*CCR5* autocrine signaling additionally drives monocyte chemotaxis and recruitment^120,121^. Monocytes were further predicted to send multiple signals to DCs, including *IL18*, which promotes DC maturation^122^ and *TNF*-*TNFRSF21*, which can induce apoptosis and inflammatory DC maturation as well as suppression of interferons^123,124^. Several NRP1-engaging interactions were also predicted in DCs; *NRP1* serves as a co-receptor on DCs that modulates both angiogenic and immune signaling, and its engagement by multiple ligands from monocytes and other DCs suggests active remodeling of the DC compartment during early infection^125^. Inhibitory interactions were also present early: *NECTIN2* on monocytes engaging *CD96* on CD8 T cells suppresses cytotoxic T cell function^126^, and *LILRB4*-*LAIR1* signaling from DCs to monocytes engages two inhibitory receptors simultaneously, indicating early onset of inhibitory feedback even during peak immune activation^127^. In contrast, late infection was dominated by broad TGF-β signaling. Both NK cells and CD8 T cells were predicted to secrete *TGFB1* engaging *TGFBR1* and *TGFBR2* across multiple target cell types including CD8 T cells, DCs, and B cells, as well as *TGFB1*-*ITGB1* interactions within NK cells which activates LAP-TGFβ. The convergence of TGF-β signaling from multiple cytotoxic effector populations onto diverse target cell types is a hallmark of immune exhaustion in chronic infection, suppressing DC antigen presentation, impairing effector CD8 T cell function, and dampening B cell responses^128^.

Taken together, these data indicate that innate inflammation dominates early infection overall; however, broad neutralizers exhibit evidence of early exhaustion programming through interferon-driven inhibitory feedback and MDSC-promoting signals, whereas narrow neutralizers preferentially engage NF-κB-driven DC activation and Th1-polarizing circuitry, suggesting that higher viral loads in broad neutralizers may drive greater immune activation with concurrent negative regulation. Immune exhaustion is highlighted in late infection with widespread TGF-β signaling across multiple cell types; exhaustion and immune checkpoint interactions were predominantly reflected in broad neutralizers, whereas narrow neutralizers maintained antiviral IFNγ-driven DC activation, underscoring a fundamental divergence of antiviral immunity in narrow neutralizers and pathologic inflammation in broad neutralizers as infection progresses.

## Discussion

Here we present the largest single-cell, multi-omic characterization of untreated HIV infection to date and the first single-cell-level analysis of immune correlates of antibody breadth in either early or chronic HIV infection. This analysis leverages a longitudinal cohort of Kenyan women living with HIV, whose monthly clinic visits enable rare profiling of early infection, allowing an understanding of immune correlates that precede breadth. Broad neutralizers had higher viral load accompanied by a steeper decline in CD4 T cells; this elevated antigen burden and disease severity is accompanied by highly activated immune cells in early infection, but progressive exhaustion in late infection. Narrow neutralizers exhibited lower viral loads and preserved immune function (Fig. 8D). Together these findings reframe antibody breadth in the setting of HIV infection as a consequence of poorly controlled infection rather than a functional anti-viral immune response.

Broad neutralizers were characterized by system-wide activation in early infection across NK cells, CD8 T cells, CD4 T cells, DCs, and monocytes with widespread ISG expression in response to viral load. Notably, NK activation, CD4 T cell help, and monocyte inflammation were the top selected variables defining broad neutralization in early infection. In late infection, this activation drives exhaustion which emerges in NK cells, CD8 T cells and DCs. While exhaustion in cytotoxic lymphocytes may be adverse to viral control, it could dampen NK cell killing of Tfh cells that would otherwise restrict breadth^34,39,45–48^. In early infection, surviving CD4 T cells maintained higher CD40L and ICOSL expression, supporting B cell help in broad neutralizers. B cells, in turn, showed early activation and SHM in the memory compartment, although this maturation did not extend to plasmablasts and did not translate into improved viral control.

Narrow neutralizers maintained relative cellular function along with lower viral loads throughout infection. CD8 TEMs, NK cells, CD4 T cells, and monocytes retained greater responses to viral-like stimulation, with NK expression of perforin and myeloid migration as top selected variables. NK cells from narrow neutralizers sustained cytotoxic capacity and expression of CXCR5 which may enable migration to lymph nodes and elimination of Tfh cells^85–87^. This could both suppress the germinal center activity required for neutralization breadth and contribute to the elimination of HIV-infected cells, which is reflected in lower viral loads^34–38,51^. Although functional cellular immunity was present, narrow neutralizers were also enriched for exhausted, atypical *NKG7*+ B cells (NKB cells), which are aberrant, chronically activated B cells identified in SIV infection^84^. NKG7 expression in B cells was the top predictor of narrow neutralization in our variable selection model which reveals that B cell-intrinsic dysfunction accompanies low antigen load in narrow neutralizers.

The tradeoff between function and exhaustion was most pronounced in CD8 T cells which are directly linked to control of HIV viremia. Narrow neutralizers, who maintain lower viral loads, are associated with progenitor exhausted T cells (which retain proliferative and cytotoxic capacity and respond to immunotherapy). Broad neutralizers, with higher viral loads, are associated with terminally exhausted T cells (which are permanently anergic)^95^. This mirrors the functional CD8 T cells associated with elite controllers (EC) (PLWH who suppress viremia in the absence of ART) and post-intervention controllers (PLWH who suppress viremia after immunological intervention and cessation of ART), suggesting that preservation of CD8 T cell function suppresses HIV replication even in those who do not fully control the virus^63–69^. Recent work shows that anti-PD-1 therapy reduced HIV reservoirs in PLWH and cancer who preserved highly functional CD8 T cells, giving mechanistic evidence that CD8 T cell function is a determinant of disease control^64^.

Underlying this divergence in CD8 T cell exhaustion state are differences in monocyte-CD8 T cell interactions. Monocytes are early and strong responders to viral infection; however, under chronic inflammation, MDSCs with functional suppressive capacity can arise to counter inflammation^129^. MDSCs can directly drive CD8 T cell exhaustion observed in broad neutralizers and are associated with poor response to immune checkpoint inhibitors^130–132^. In contrast, narrow neutralizers had early Treg accumulation, which may dampen inflammation and preserve CD8 T cell and other immune function. Differences in exhaustion emerged in early HIV infection, indicating that CD8 T cell exhaustion could be relevant in PLWH who receive treatment soon after acquisition. This viral load driven CD8 T cell exhaustion state appears to be predictive of response to immunotherapeutic strategies aimed at HIV cure.

Cellular programs correlated with activation followed by exhaustion in broad neutralizers were consistent across the immune system. NK cells, CD4 T cells, B cells, CD8 T cells, monocytes, and DCs were enriched for AP-1, NF-κB and/or interferon-response pathways in broad neutralizers in early infection followed by enrichment for metabolic pathways in late infection, reflecting the metabolic reprogramming in exhausted cells. In contrast, narrow neutralizers exhibited enrichment of AP-1 and NF-κB related pathways predominantly in late infection, suggesting delayed inflammation correlated with retained immune function and more controlled viral load. NF-κB can be activated by TNF signaling which was prevalent in broad neutralizers and drives type I interferon, MHC I expression, AID mediated class switching, and SHM during HIV infection^133^. However, NF-κB and AP-1 also synergistically promote latency reversal and HIV transcription, and HIV’s nef protein negatively regulates NF-κB, indicating its importance in immunity to HIV^134,135^. This tension between anti-viral immunity and viral transcription may explain why broad neutralizers, despite exhibiting greater early immune activation, have concurrent higher viral load and ultimately develop greater systemic immune dysfunction. The metabolic shift observed in late infection across multiple cell types in broad neutralizers is consistent with this interpretation; cells undergo metabolic reprogramming as a hallmark of exhaustion-associated dysfunction observed in both chronic viral infection and cancer^136,137^. These data suggest that viral load is not merely a correlate of antibody breadth but a mechanistic driver of the transcriptional and functional immune remodeling that enables breadth.

Although our study offers an unprecedented look into untreated HIV infection, several limitations of this study warrant consideration. Our cohort consists exclusively of Kenyan women who engage in sex work. While this cohort has been indispensable for studying natural HIV immunity, generalizability to other populations, sexes, and clades is not guaranteed. The neutralization panel described here does not identify the presence of monoclonal bnAbs but instead reflects the natural spectrum of breadth rather than an absolute threshold. Within our cohort, broad neutralizers were older and had a greater number of previous pregnancies. While no pregnancies occurred during the study period, both factors could independently shape immune set point. Adjusting for age and number of pregnancies did not attenuate most associations and strengthened several, indicating negative confounding of these immune features. More broadly, bootstrapped resampling of effect sizes for all Stabl-selected features confirmed that associations were robust to donor-driven effects. Our longitudinal design captures immune trajectories across early and late infection; however, we can only draw conclusions from correlations between immune states and neutralization breadth due to limitations of profiling. While the role of MDSC-driven CD8 T cell exhaustion will require direct experimental interrogation, NK cell-mediated suppression of Tfh cells^34,51^, the role of CD8 T cell exhaustion state^138^, and Treg accumulation in narrow neutralizers^14^ are each supported by prior work. Finally, all participants in this cohort were untreated, and ART initiation could alter the immune trajectories described here, though the early emergence of differing exhaustion programs suggests that treatment timing would be an important factor in these dynamics.

These findings paint a picture of two divergent immunological trajectories shaped early in HIV infection by the degree of viral control. Our data suggest a feed-forward loop in which initial differences in viral load are reinforced over time by their downstream effects on immune function. In broad neutralizers, high antigen burden drives acute immune activation and early B cell activity that precedes antibody breadth, but ultimately this early peak in activation drives progressive exhaustion, MDSC-mediated suppression, and loss of viral control. Narrow neutralizers preserve effector function and more effective viral suppression, but lack stimulation to drive antibody maturation toward broad neutralization. This framing has critical implications for HIV vaccine and cure strategies. The conditions of a favorable immune response (low viral load) in HIV-infected individuals requires preserved cellular immunity and is at odds with those that favor antibody breadth. Exhaustion of cellular immunity will be both a prime target and a key barrier in HIV cure approaches, while recapitulating the sustained inflammation required to develop broad antibodies will remain a central challenge for vaccination.

## Methods

### Study Design

Samples were obtained from the Mombasa Cohort, a long-term open cohort study of women who engage in sex work located in Mombasa, Kenya. Established in 1993, the cohort enrolls HIV-negative women who engage in sex work and tests monthly for HIV acquisition enabling accurate assessment of HIV acquisition date and profiling of acute HIV infection. The timing of the first infection is defined by both HIV serology and HIV RNA testing. If they acquire HIV, women are asked to return to donate PBMCs and plasma quarterly. All women in the cohort are treated for HIV infection according to Kenyan national guidelines. Starting in 2004, ART was initially only available to those with a CD4 count <200 cells/μL or an AIDS-defining illness. CD4 T cell count thresholds for ART initiation increased over time through 2015, and the recommendation to treat all PLWH was implemented in 2016. The present study uses samples collected before 2012 from ART-naive PLWH and represents longitudinal samples of untreated HIV infection.

Individuals were selected for profiling if they had matched PBMC samples from early (14 - 67d post infection) and late (2.6 - 7.6y post infection) infection timepoints. 27 selected individuals were profiled by plasma neutralization assays performed on late infection timepoint. No individuals were receiving ART treatment, TB treatment, or met the clinical definition of AIDS during selected sample timepoints. ScRNA-seq and CyTOF profiling was performed on n = 16 early infection and n = 18 late infection PBMC samples due to quality-based sample drop out (Fig. S1A). Additional control samples from HIV-negative individuals (n = 6) in the same cohort were also included. Blood was collected in heparin tubes; plasma was isolated by centrifugation and PBMCs were isolated by ficoll gradient. Plasma was kept at −80C and PBMCs at −170C in liquid nitrogen for long term storage. Verbal or written informed consent was obtained from all patients. The ethical review committees of the University of Nairobi, the University of Washington, and the Fred Hutchinson Cancer Research Center approved this study.

### Sex as a Biological Variable

The Mombasa female sex worker cohort includes only females and to our knowledge, similar samples are not available from untreated male subjects, nor is it ethical to collect such samples at this time as ART is the standard of care globally. We therefore conducted a single sex study.

### Pseudotyped Virus Production

HIV envelope (env) pseudotyped viral particles were produced in 293T cells (ATCC) using co-transfection of cloned viral envelope sequences and a full-length subtype A proviral clone with partial deletion in envelope (Q23Δenv) as previously described^138^ using Fugene 6 transfection reagent (Promega; E2691). Two million cells were seeded in D10 (DMEM (Life Technologies; 11885-092) + 10% FBS, 1% L-glutamine, 1% PSA, 10mM HEPES) in T75 flasks 24h before transfection. The env plasmid consists of approximately 9000 base pair pCI-neo (Promega) or pCDNA3.1-TOPO backbone with an approximately 3000 base pair env sequence. Q461.d1^139^ – GenBank ID: AF407155; SF162^139^: EU123924; Q842.d16^140^: AF407162; DU156.12^141^: DQ411852; Q435.100M.a4^138^: FJ866139; Q461.e2^138^: AF407156; Q23^141^: AF407153; Q406.70M.f3^138^: FJ866133; Q259.d2.d26^138^: AF407153; Q769.b9^142^: AF407157. All plasmids were filtered through a 0.22μM filter for sterility before use. In a final volume of 188 μL, 2.67 μg of Q23Δenv and 1.33μg of envelope plasmid were mixed with serum-free DMEM. 12 μl of Fugene6 was added dropwise and incubated at room temperature (RT) for 15m to form transfection complexes. The 200μL transfection mix was then added dropwise to plated 293T cells. 24h after transfection, culture media was replaced. Viral supernatants were harvested 72h after transfection by 0.45μM filtration. Viral stocks were aliquoted and stored at −80C until further use (no less than 48h).

### Pseudovirus Neutralization Assay

TZM-bl, a genetically engineered HeLa cell line that expresses CD4, CXCR4, and CCR5 and contains Tat-responsive reporter genes for firefly luciferase (Luc) and *Escherichia coli* β-galactosidase under regulatory control of an HIV-1 long terminal repeat, was used as target cells^143^. TZM-bl cells were thawed in D10 and passaged at least twice before use. Patient sera were inactivated at 56C for 1h before use. Sera diluted 1:50 was filtered through a .22μM spinX filter (Corning; 8160). Serial dilutions (1:2) of sera up to 1:1600 was performed in 96-well black walled, clear bottom plates (Thermo Fisher; 165305) in 25μL final volume. 25μL of pseudovirus was added at a titer such that virus-only wells would achieve a luminescence of approximately 100,000 RLU on Promega Glomax plate reader (Promega; GM3000). Virus-serum mixtures were incubated for 1h at 37C. TZM-bl cells were harvested using trypsin (Thermo; 25200072). After incubation 10,000 cells (50μL) per well were added; 48h later, all media was removed from plate by vacuum aspiration and cells were lysed by addition of 50μL of Britelite Plus assay readout solution (Revvity; 6066766) and 50μL of 1X PBS. Luminescence was measured using Promega Glomax after 30s shaking. Background luminescence (cell only wells) was averaged and subtracted from all wells. Percent neutralization was calculated by comparing to 100% infection (virus only wells). The reciprocal of the highest dilution to obtain <50% infection was used to calculate average half maximal neutralization titer (NT50). NT50 above 100 (limit of detection) was counted as neutralization.

### Patient Sample Thawing

All work with primary cells was performed with low retention pipette tips. PBMCs were thawed at 37C and washed with RP10 (RPMI (Gibco;11875093), 10% FBS) with 20μL of benzonase (Millipore Sigma; 70664-3). Samples were counted to determine viability and dead cell removal (Milteny; 130-090-101) was performed on samples with <70% viability according to manufacturer’s recommendation. Samples were spun down for 5m at 300g, resuspended in RP10, and passed through 70μM MACS Smart Strainer (Miltenyi; 130-098-458) to create a single cell suspension. Cells were then counted, spun down and resuspended at 2e6cell/mL and 50μL were transferred to DNA LoBind tubes on ice (Eppendorf; 022431021) for scRNA-seq.

### 10X 5’ scRNA-seq

Cells were counted in duplicate and RP10 was added to adjust the concentration to 1e6 cells/mL (total cells, not live cells). 10,000 cells were loaded on Chromium Next GEM Chip K according to recommendations. Cells and gel beads were partitioned using the 10X Genomics Chromium platform using the Chromium Next GEM Single Cell 5′ Reagent Kits v2 (10X; CG000331). After RNA capturing, cDNA synthesis, and V(D)J amplification, libraries were constructed using Dual Index Kit TT Set A (10X: 1000215) according to manufacturer’s recommendations. Quality control was performed on Agilent Tapestation using HSD5000 kit to ensure no primer peaks were detected. Libraries were sequenced on NovaSeq X instrument to achieve a target sequencing depth of 20,000 reads per cell for GEX and 5,000 reads per cell for V(D)J.

### Alignment, Quality Control, and Pre-processing of scRNA-seq data

Cell Ranger V7.2.0 was used to align the GEX and VDJ FASTQ reads to the human genome using the multi pipeline to improve V(D)J alignment. The resulting gene expression matrices were used to create a Seurat object (V5.3.0) in R (V4.5.1). Cells that had <500 or >5,000 UMIs, cells with <1,000 or >12,000 total counts, cells with >20% gene expression from mitochondrial cells, cells with <2% or >16% RPS expression, cells with < 2% or >20% RPL expression, and cells with >1% red blood cell gene expression were considered low quality and were removed. Doublets were identified using scds^144^ and cells with hybrid score >1 were removed; this was consistent with the ∼8% doublet rate expected. Data were normalized using SCTransform() with percent mitochondrial genes, percent ribosomal features, total counts, and total UMI as confounding variables. TCR and BCR genes removed from variable features prior to dimensionality reduction to avoid clustering artifacts^145^. Principal component (PC) analysis (PCA) was performed using the 3,000 most highly variable genes. The Harmony algorithm was applied to correct for donor-level batch effects; the first 50 dimensions were used for graph-based clustering and UMAP visualization.

### V(D)J Gene Alignment

V(D)J Analysis was performed using the Immcantation Framework. V(D)J contig sequences generated by CellRanger were used to assign V, D, and J gene segments using Change-O^146^ package and IgBLAST^147^ against the IMGT human immunoglobulin germline reference database. Cells with multiple heavy chains or without heavy chains were removed. Clonal groups were defined using hierarchicalClones() function from the SCOPer package, employing findThreashold(). Germline sequences were reconstructed for each clonal family using CreateGermlines() with the IGMT germline reference dataset. Somatic hypermutation frequencies were calculated using the SHazaM package observedMutations() function. BCR features were assigned to cells using cell barcodes.

### Mass Cytometry Staining Cocktail

Surface and intracellular antibodies were either pre-conjugated from Fluidigm/Standard BioTools or conjugated using Fluidigm Maxpar X8 labeling kits or the Ionpath MIBItag conjugation kit (for 157Gd only) (Table S9). Unconjugated Maxpar Ready antibodies were used when possible. Antibody-metal conjugations were performed according to manufacturers’ instructions with the following deviations for the Maxpar X8 labeling kits: 30kDa filters were used instead of 50kDa filters and metal-conjugated antibody was recovered in W buffer. Concentration was determined using Qubit 4 Fluorometer (Thermo Fisher) per manufacturer instructions. Conjugated antibodies were then titrated, pre-combined into either surface or intracellular staining cocktails (Table S9). Cocktails were filtered through a 0.22μM polyethersulfone syringe filter and frozen at - 80C for long-term storage.

### Mass Cytometry

After the 10X chip was loaded, remaining cells were split for a stimulated and unstimulated condition. Up to 2e6 cells per condition were resuspended in 200μL of complete RP10 (RPMI (Gibco;11875093), 10% FBS, 1% PSA, 1% L-glutamine, 10mM HEPES, 1% NEAA, 1% Sodium Pyruvate) and rested overnight in 96-well U-bottom plates. After overnight rest, stimulated cells were resuspended in 4μg/mL R848 (Invivogen; tlrl-r848) and 12.5μg/mL Poly(I:C) (Sigma; I3036-2MG) for 5h total. After 1.5h, Brefeldin A (eBioscience; 00-4506-51), Monensin (eBioscience;00-4501-51), and anti-CD107a APC were added to all wells. After 3 hours, 0.25X PMA/I (eBiosceinces; 00-4970-93) was added to stimulated wells. This stimulation was described in Hamlin et al. and designed to optimize cytokine production in PBMCs without cell loss in healthy donors^148^.

After stimulation concluded, cells from stimulated and unstimulated conditions were resuspended in CyFACS (1XPBS (Rockland; MB-008), 0.1% BSA, 2 mM EDTA, 0.05% sodium azide prepared in single-use plastic). Samples were barcoded as using a six-choose-three scheme with an anti-b2m antibody conjugated to 194Pt, 195Pt, 196Pt, 198P, 113In and 115In, enabling live-cell barcoding of up to 20 samples^149^. After 30m at RT, cells were washed in CyFACS, resuspended in CyPBS (1XPBS prepared in single-use plastic), and pooled in a 5mL polypropylene FACS tube for staining. All spins before fixing were performed at 500g for 5m. (Ethylenediamine)palladium(II) chloride (PdCl) stored in DMSO was used for viability staining as previously described^149,150^. Pooled samples were stained with PdCl viability reagent (final concentration of 500nM in CyPBS) for 5m at RT, then quenched with an equal volume of FBS and washed twice in cyFACS. Cells were resuspended in 100μL human Fc block (Biolegend; 422302) in 2mL of CyFACS for 10m at RT. Cells were then spun down and resuspended in the surface antibody cocktail for 30m at 4C. Cells were washed twice in CyFACS, then fixed in 2% PFA for 20m at RT, protected from light. After fixing, cells were washed twice in 1X permeabilization buffer (eBioscience; 00-8333-56); all spins after fixing were performed at 800g for 5m. Cells were resuspended in the intracellular antibody cocktail for 45m at 4C. Cells were washed twice with permeabilization buffer and twice with CyFACS. Cells were then resuspended in 40nM Cell ID Intercalator-Ir (Standard Biotools; 201192B) in 2% PFA. Cells were kept at 4C, protected from light until acquisition on CyTOF Helios. Immediately before acquisition, cells were washed once and resuspended in filtered MilliQ water with 0.1x EQ Four Element Calibration Beads (Standard BioTools; 201078) and filtered through a 35μm nylon mesh into polystyrene tubes (Falcon; 352235) for sample acquisition.

### Preprocessing of Mass Cytometry Data

Flow cytometry standard (FCS) files were normalized and debarcoded using the Premessa Package (V0.3.4) as previously described^151^. FlowJo V10.10.0 was used to manually gate out EQ beads, low quality cells, doublets, debris, and dead cells using Standard Biotools clean up strategy^152^ (Fig. S9A). Cell types were manually gated as shown in S8B and S9A to further exclude low quality cells. FCS files were used to create a Seurat object using the Seurat and FlowCore packages with manual cell type labels. Data was arcsinh-transformed data (cofactor = 5) and UMAP embeddings were calculated using the uwot package (V0.2.2) excluding IFN-G and TNF-A to avoid stimulation-based clustering effects. All box plots depicting CyTOF data represent transformed per-sample mean signal intensity (transformed MSI).

### Annotation of Cell Types in scRNA-seq Data

Broad cell types were assigned using three independent methods. First, the Seurat implementation of a weighted nearest neighbors strategy was used to transfer cell labels from a large multi-modal reference data set^153,154^ using FindTransferAnchors() to identify anchors between the reference and query dataset and MapQuery() to transfer cell type labels. Second, SCimilarity, a pre-trained deep-metric-learning foundation model was used to transfer the most similar cell type labels from the annotation model V1^155^. Third, Single R was used for correlation-based reference assignment of cell types using bulk RNA-seq references^92^. Here, the ImmunoStatesMatrix from the Metaintegrator package was used as a reference which contains 20 purified immune cell populations. For broad cell type annotations (e.g. CD4 T, NK etc.), agreement of 2/3 methods was used to assign cell types. Remaining cells were re-clustered and manual annotation of cellular identity was performed by finding DEGs for each cluster using Seurat’s implementation of the Wilcoxon rank-sum test (FindAllMarkers()) and comparing those markers with known cell type-specific genes. Where relevant, fine cell type annotations (i.e. B Naive, B Memory etc.) were assigned by the agreement of 2/3 methods where possible; if fine cell types were not present in two or more methods, manual annotation was used.

### Differentially Expressed Genes in scRNA-seq Data

Differentially expressed genes (DEGs) were identified by Seurat’s implementation of the Wilcoxon rank-sum test (FindMarkers()) with logfc.threshold = 0.25, min.cells.feature = 100, min.pct = 0.05.)

### Blood Transcriptional Module Analysis

Pathway analysis using blood transcriptional modules (BTMs) was performed as previously described ^75^. Upregulated DEGs were used as input. Only modules with >1 gene represented and adjusted p <.05 were displayed. Up to the top 10 BTMs were visualized.

### Feature Selection

To identify top features within each data modality that were predictive of breadth, Stabl (V1.0.0) was applied independently to each data modality (RNA, protein, BCR, and clinical) using reticulate (V1.43.0) in R (Table S4)^98^. Stabl is a supervised machine learning framework that couples noise injection via Model-X knockoffs with a data-driven signal-to-noise reliability threshold to identify sparse, reliable feature sets within a multivariable predictive modeling architecture. Elastic net (EN) regression was used to capture correlated variables; briefly, Stabl fits EN models across repeated subsamples of the data to estimate each feature’s selection frequency, then injects artificial knockoff features to derive a false discovery proportion surrogate. The reliability threshold is defined as the selection frequency that minimizes false discovery proportion. This procedure was applied independently to each modality, allowing a modality-specific reliability threshold to be computed for each dataset. To characterize individual associations between selected features and breadth group, post-hoc univariable Firth penalized logistic regression was performed for each selected feature within each modality to estimate the log-odds coefficient (Table S5). Firth penalization was used in place of standard logistic regression to obtain bias-corrected coefficient estimates appropriate for the small sample size, where standard maximum likelihood estimation is susceptible to separation.

### Confounder Analysis

To assess whether Stabl-selected features associated with breadth were confounded by age, gravidity, or viral factors we performed Firth penalized logistic regression with covariate adjustments for selected feature sets. The unadjusted model (M0: Breadth∼feature), a model adjusted for demographic factors (M1: Breadth ∼ feature + age + pregnancies at enrollment), and a fully adjusted model (M2: M1: Breadth ∼ feature + age + pregnancies at enrollment + plasma viral load + CD4 T cell count (late timepoint only) were compared to determine the degree of confounding (quantified as percent moderation: β_Mx − β_M0) / |β_M0| × 100). Positive values indicate that the unadjusted association was suppressed by the covariate (negative confounding) and negative values indicate attenuation of the association after adjustment

### Bootstrap Stability Analysis

To address robustness of Stabl-selected features to donor sampling, we performed stratified bootstrap resampling (B=1000) of Hedge’s g for each selected feature. Broad and narrow neutralizers were sampled independently with replacement, and the 95% confidence interval was estimated by the 2.5 and 97.5th percentiles of the bootstrap distribution.

### Gene Module Scoring

The Seurat function AddModuleScore() was used to score single cells by expression of a list of genes (Table S7). This function calculates a module score by comparing the expression level of an individual query gene to other randomly selected control genes expressed at similar levels to the query genes and is therefore robust to scoring modules containing both lowly and highly expressed genes, as well as to scoring cells with different sequencing depths.

### MultiNicheNet Analysis

MulitnichenetR (V1.0.1)^156^ was used to identify differentially expressed active receptor-ligand interactions between broad and narrow neutralizers. Top 25 receptor-ligand interactions were identified using logFC_threashold = .5, p_val_threashold = .05, fraction_cutoff = .05 (Table S8). These receptor-ligand interactions were manually verified in the literature. Interactions between intracellular receptors erroneously identified by nichenet were colored transparent (i.e. interactions with HLA-DO).

### Data Visualization

All data analysis and visualizations were performed in the open-source software R (V4.2.2)^157,158^. The R package Seurat (V5.3.0) was used to generate UMAP projections. ComplexHeatmap (V2.24.0) and ggchicklet2 (V0.7.0) were used for heatmaps. MultinichenetR was used to create circos plots. Colors for plots were generated using paleteer. Custom ggplot2 functions were used for all other plots.

### Data availability

FCS files (mass cytometry) with de-identified metadata supporting this publication will be available upon publication at Cytobank Community. Data from scRNA-seq will be available upon publication at Gene Expression Omnibus.

### Code Availability

A Github repository for all code to reproduce analyses and figures will be available upon publication at https://github.com/BlishLab/HIV_breadth

### Statistical Analysis

Comparison of means was performed with the Wilcoxon rank sum test (two-sided). P-value adjustment was performed by bonferroni correction where applicable. The threshold for statistical significance of differential gene expression in scRNA-seq data for each cell type was determined as adjusted P < 0.05.

## Acknowledgements

We thank Drs. Peter S Kim, Payton Weidenbacher, Theodora Bruun, and Duo Xu for the gift of DU156.12 env plasmid and assistance with pseudovirus neutralization assay protocol. We thank Drs. Rebecca Hamlin and Leslie Chan for their development of and assistance with CyTOF stimulation protocol. We thank Dr. Brooks Benard for his assistance with data visualization. Q461.e2 and SF162 env plasmids were obtained through the NIH HIV Reagent Program, Division of AIDS, NIAID, NIH: Human Immunodeficiency Virus 1 (HIV-1) gp160 Expression Vector (Q461ENVe2), ARP-10460, contributed by Dr. Julie Overbaugh; SF162 gp160 Expression Vector, ARP-10463, contributed by Dr. Leonidas Stamatatos and Dr. Cecilia Cheng-Mayer. This work was supported, in whole or in part, by the Bill & Melinda Gates Foundation OPP1113682. Under the grant conditions of the Foundation, a Creative Commons Attribution 4.0 Generic License has already been assigned to the Author Accepted Manuscript version that may arise from this submission. C.A.B. is an investigator of the Chan Zuckerberg Biohub. This work was also supported by fellowship and training support from National Institutes of Health through Grants T32 AI007290 Molecular and Cellular Immunobiology, SGF: Stanford Graduate Fellowship in Science & Engineering, and Gerald J. Lieberman Fellowship (to I.D.K.). Additional support, including to the cohort, was provided by R01 HD103571 (to J.O.).

## Supplementary Figures

**Supplementary Fig. 1:**
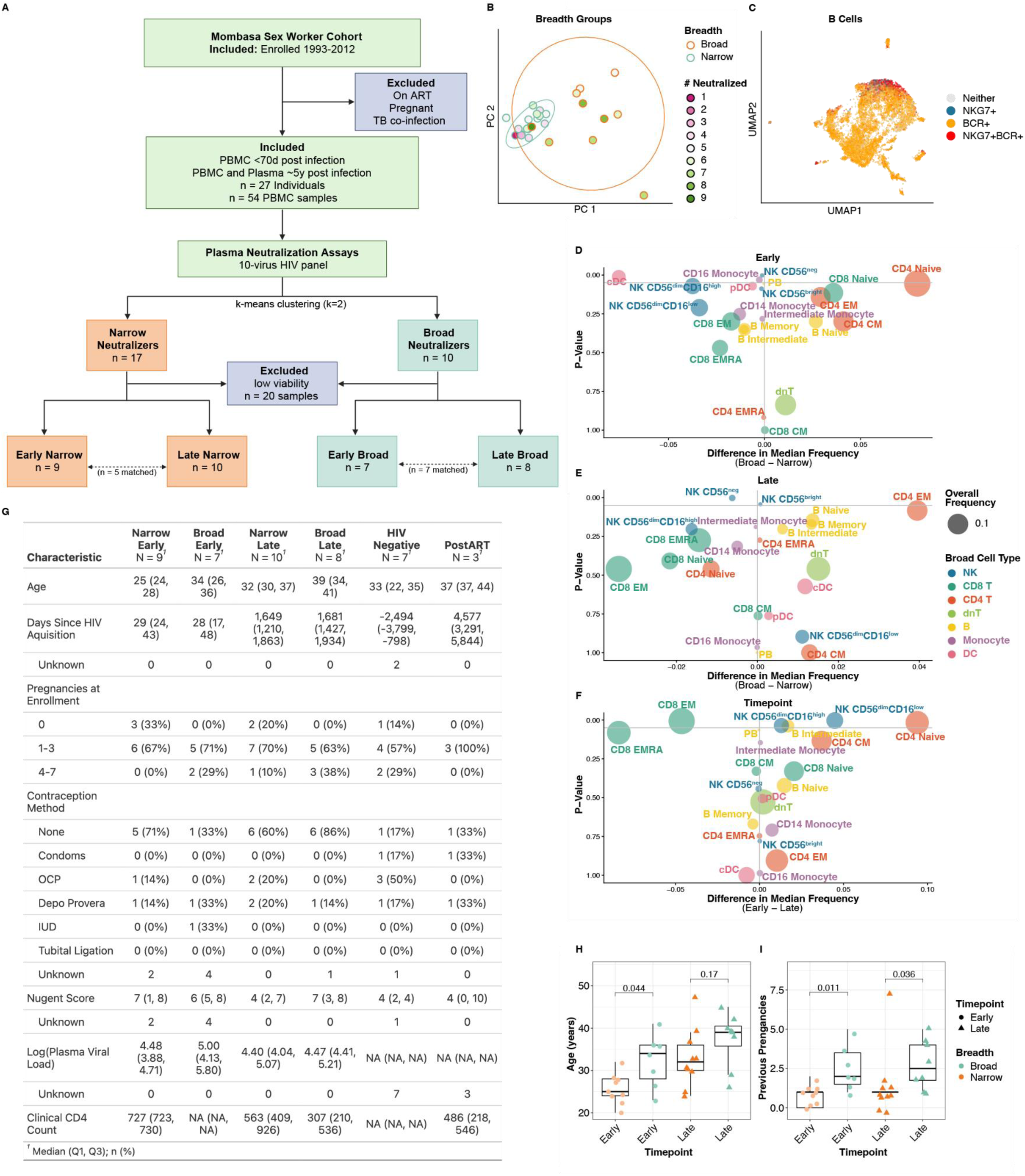
Study metadata. **A.** Sample outline for study design and data exclusion. **B.** Scatterplot of principal component (PC) 1 and 2 from PCA of NT50 across 10 HIV pseudoviruses. Each point represents one subject colored by the number of viruses neutralized. Outlines represent breadth groups defined by k-means clustering (k = 2). **C.** UMAP of B cells from scRNA-seq dataset colored by the presence of a successfully sequenced BCR and/or *NKG7* expression above zero. **D-F.** Volcano plot of median difference in fine cell type frequencies (% of PBMC) in early and late infection (**D**), broad and narrow neutralizers in early infection (**E**), and broad and narrow neutralizers in late infection (**F**). Y axis shows P-value from wilcoxon rank sum test. **G.** Clinical and demographic characteristics of study participants by breadth group and infection timepoint. NA represents missing or irrelevant data. **H.** Boxplot of participant age in each breadth group. **I.** Boxplot of pregnancies before enrollment in breadth groups. P-values by two-sided Wilcoxon rank-sum test with Bonferroni’s correction for multiple hypothesis testing. Each point represents one donor.

**Supplementary Fig. 2:**
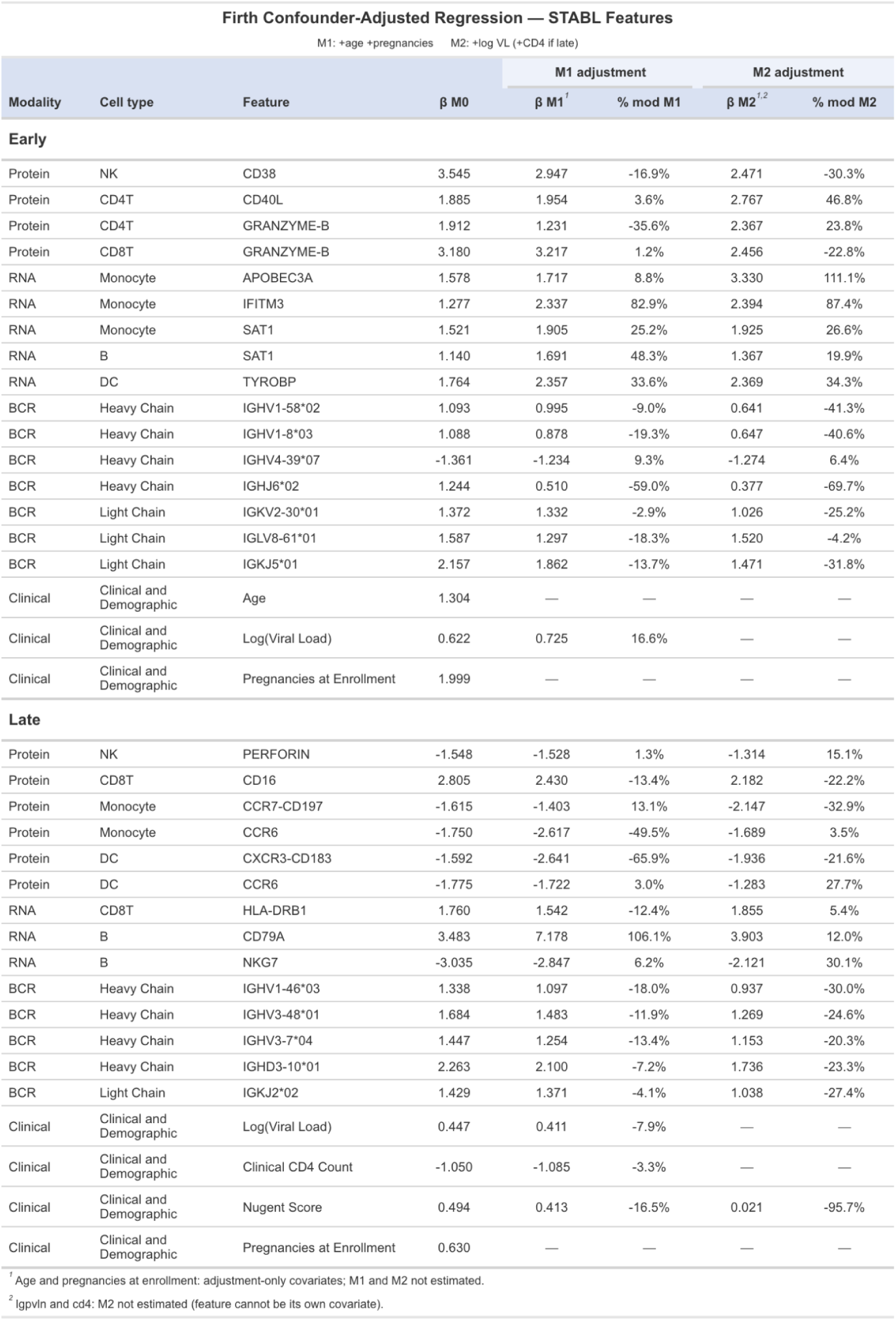
Confounder-adjusted Firth penalized regression results. Features selected by Stabl were evaluated for percent moderation after adjusting for age, pregnancies, plasma viral load, and CD4 T cell count.

**Supplementary Fig. 3:**
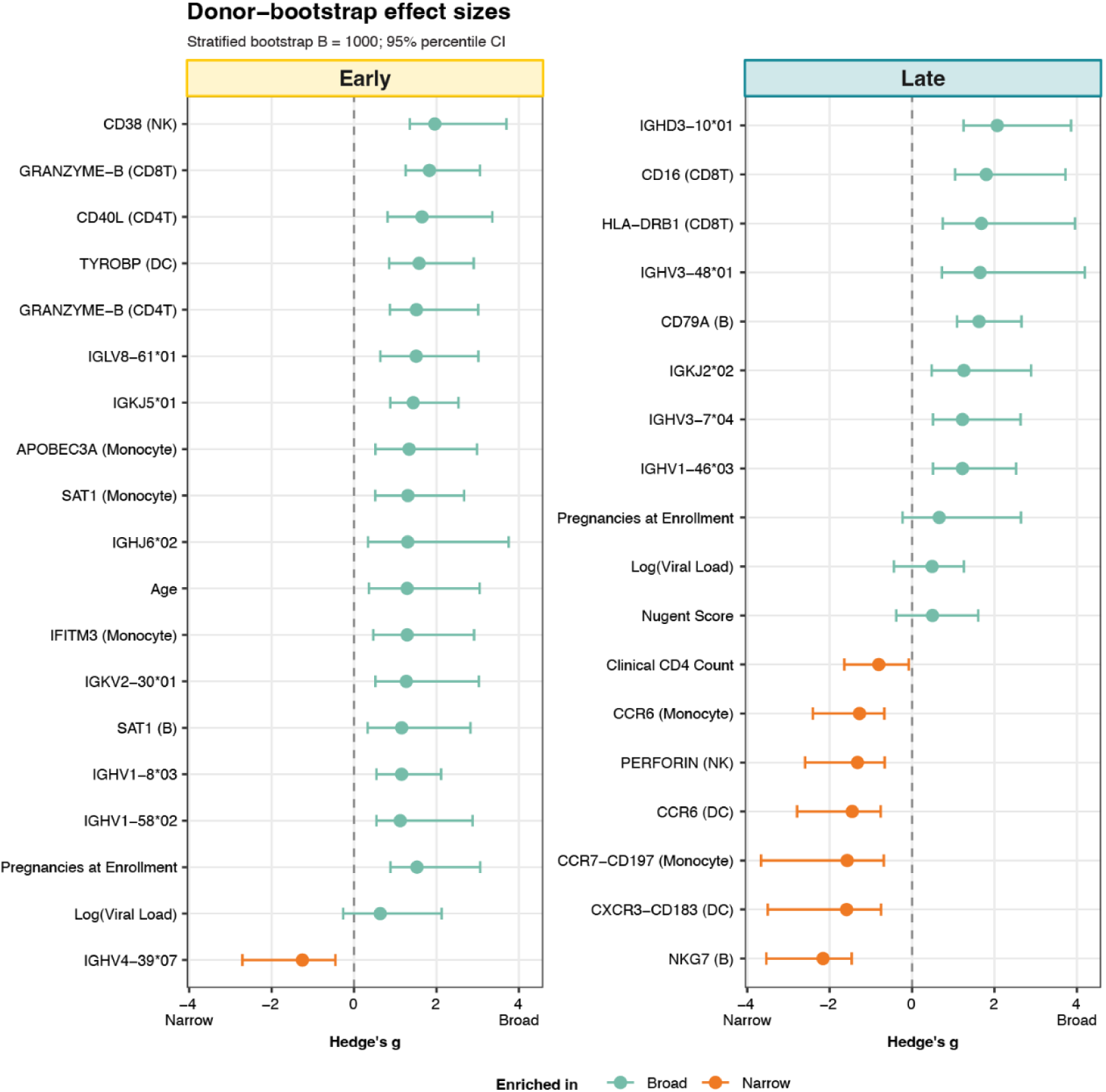
Robustness to donor bootstrapping. Hedge’s g was calculated for each Stabl-selected feature in stratified bootstrap resampling (B=1000) groups, independently resampling broad and narrow donors with replacement. Points represent the observed effect size, and error bars estimate the 95% confidence interval (CI).

**Supplementary Fig. 4:**
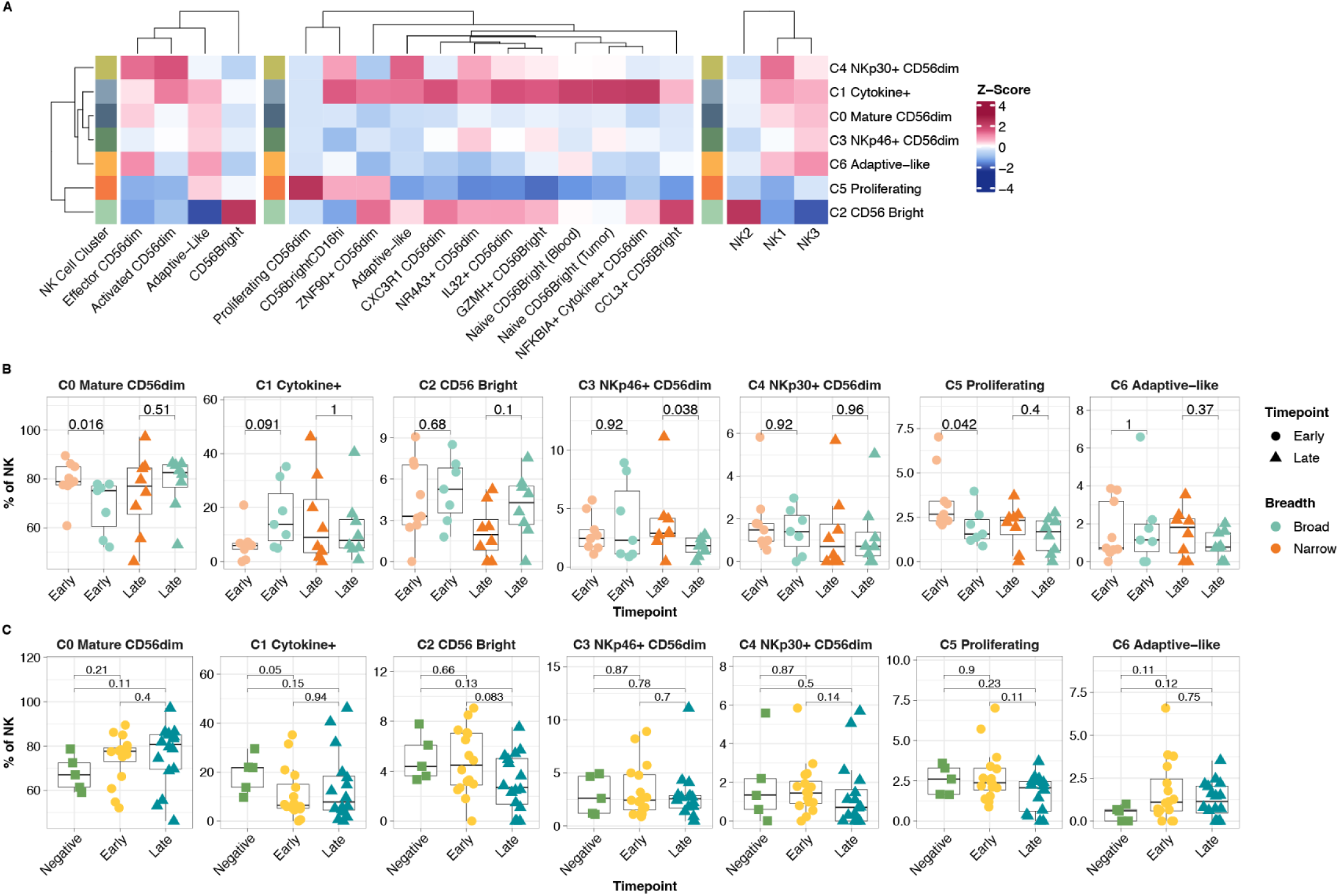
NK cell transcriptomic clusters. **A.** Heatmap of normalized expression of NK cell gene signatures from Netskar et al., Tang et al., and Rebuffet et al. in transcriptomic clusters. **B.** Boxplots of cluster frequencies in breadth groups. **C.** Boxplots of cluster frequencies in infection timepoints. P-values by two-sided Wilcoxon rank-sum test with Bonferroni’s correction for multiple hypothesis testing. Each point represents one donor.

**Supplementary Fig. 5:**
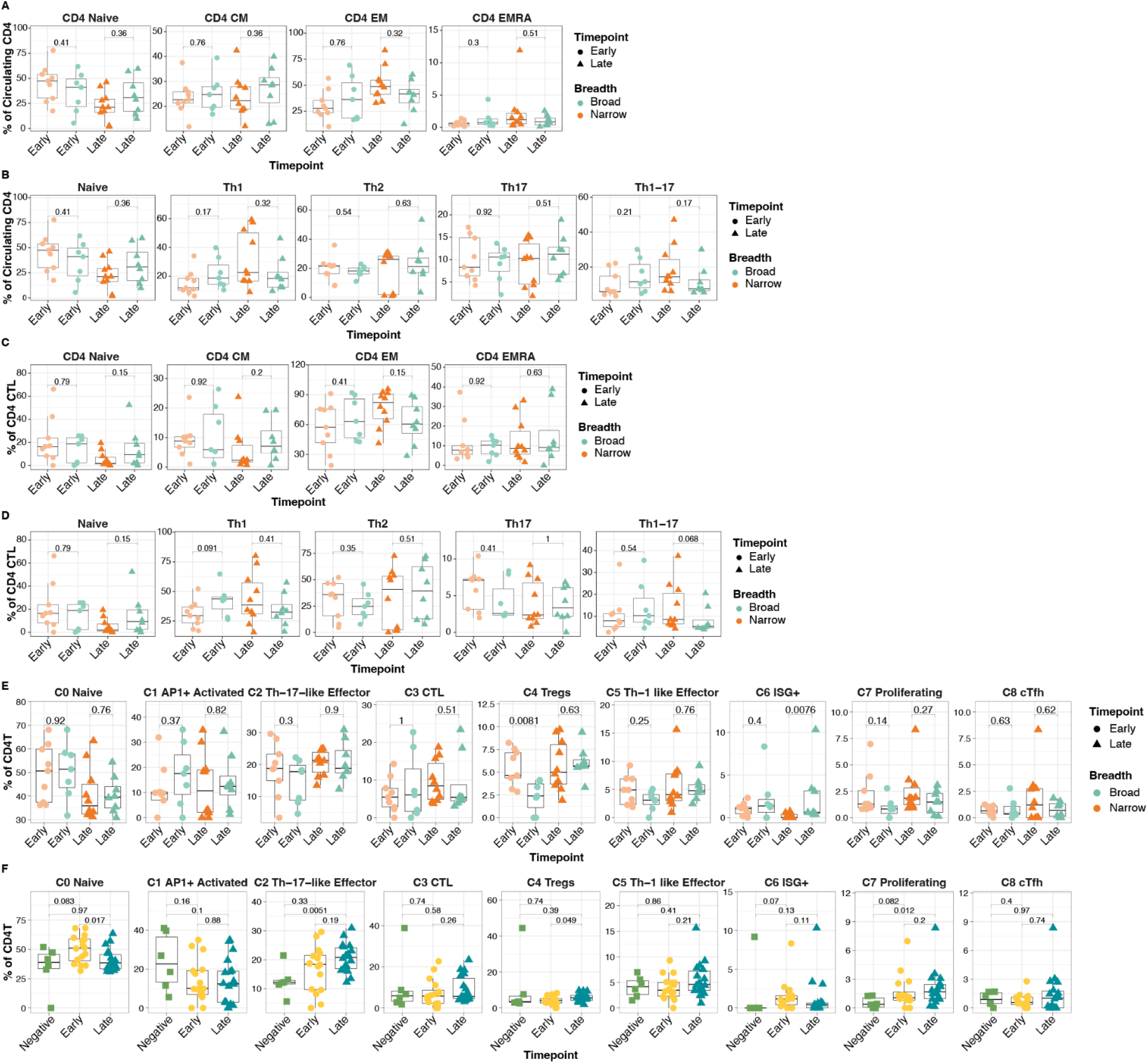
CD4 T cell phenotypes and cell transcriptomic clusters. **A.** Boxplots of frequencies of CD4 T memory types between breadth groups. **B.** Boxplots of frequencies of CD4 T helper types between breadth groups. **C.** Boxplots of frequencies of memory types among CD4 CTL between breadth groups. **D.** Boxplots of frequencies of CD4 T helper types among CD4 CTL between breadth groups. **E.** Boxplots of cluster frequencies in breadth groups. **F.** Boxplots of cluster frequencies in infection timepoints. P-values by two-sided Wilcoxon rank-sum test with Bonferroni’s correction for multiple hypothesis testing. Each point represents one donor.

**Supplementary Fig. 6:**
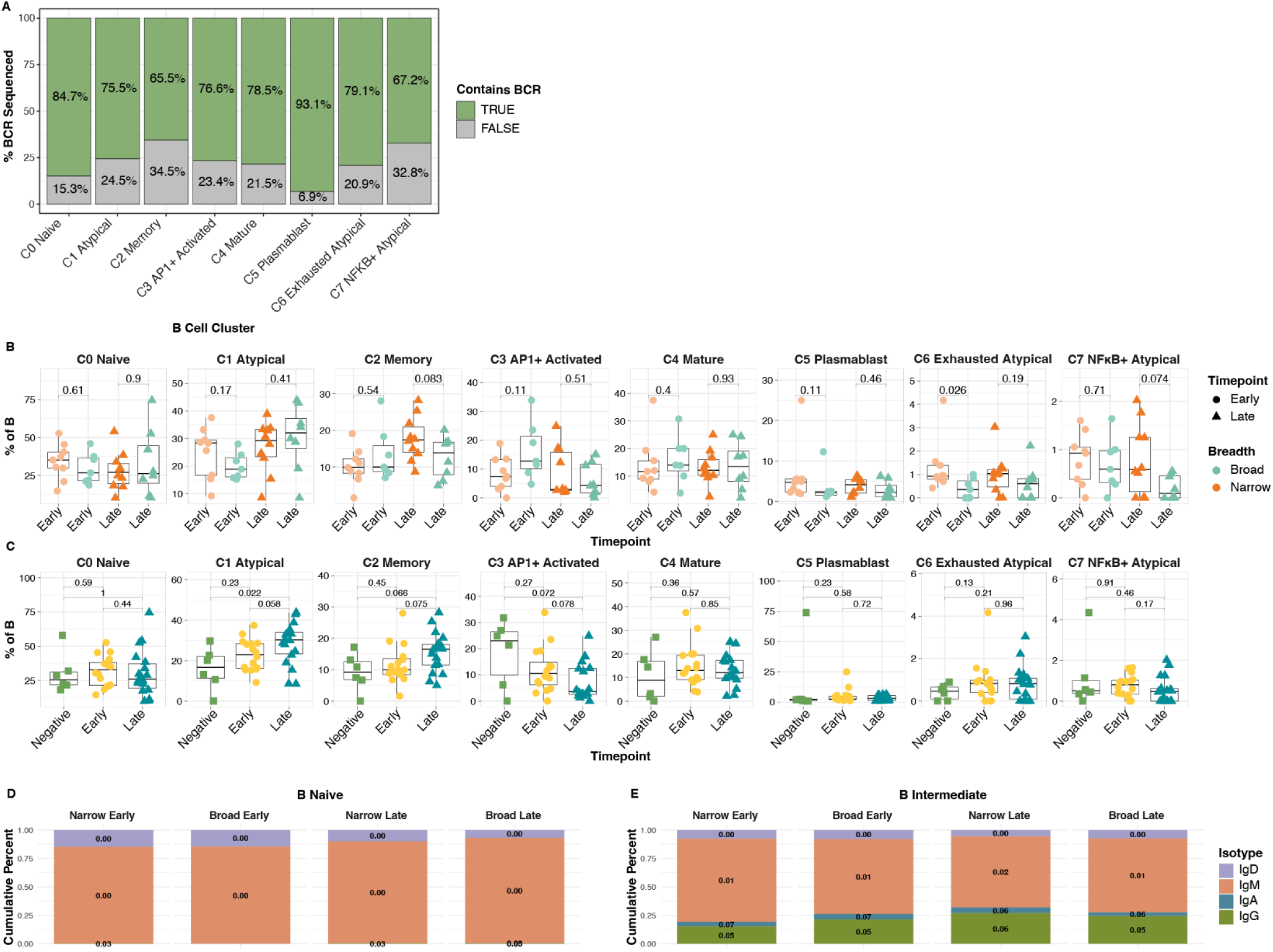
B cell V(D)J sequencing and transcriptomic clusters. **A.** Frequency of cells with BCR sequence in transcriptomic clusters. **B**. Boxplots of cluster frequencies in breadth groups. **C.** Boxplots of cluster frequencies in infection timepoints. P-values by two-sided Wilcoxon rank-sum test with Bonferroni’s correction for multiple hypothesis testing. Each point represents one donor. **D** and **E.** Cumulative bar plots of immunoglobulin heavy chain gene usage from naive B cells (**D**) and intermediate B cells (**E**) in breadth groups colored by gene with median SHM of each isotype displayed.

**Supplementary Fig. 7:**
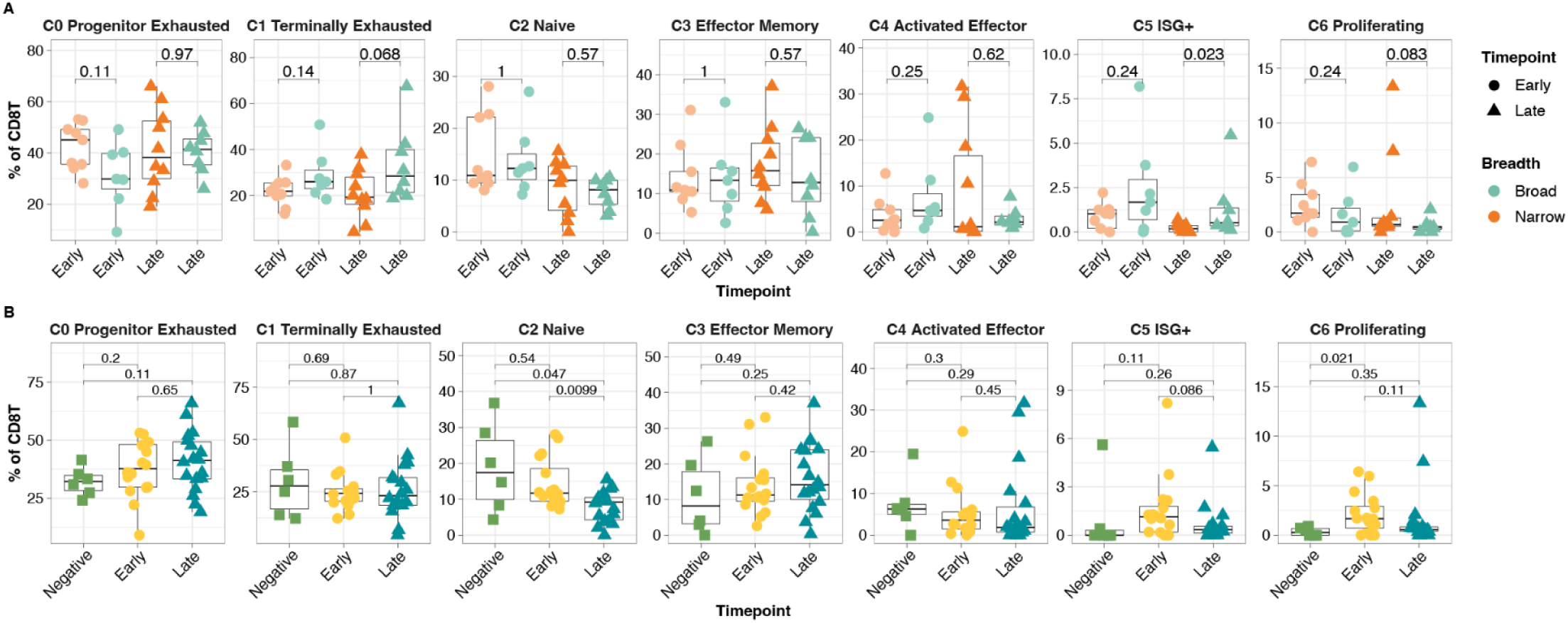
CD8 T cell transcriptomic clusters. **A.** Boxplots of cluster frequencies in breadth groups. **B.** Boxplots of cluster frequencies in infection timepoints. P-values by two-sided Wilcoxon rank-sum test with Bonferroni’s correction for multiple hypothesis testing. Each point represents one donor.

**Supplementary Fig. 8:**
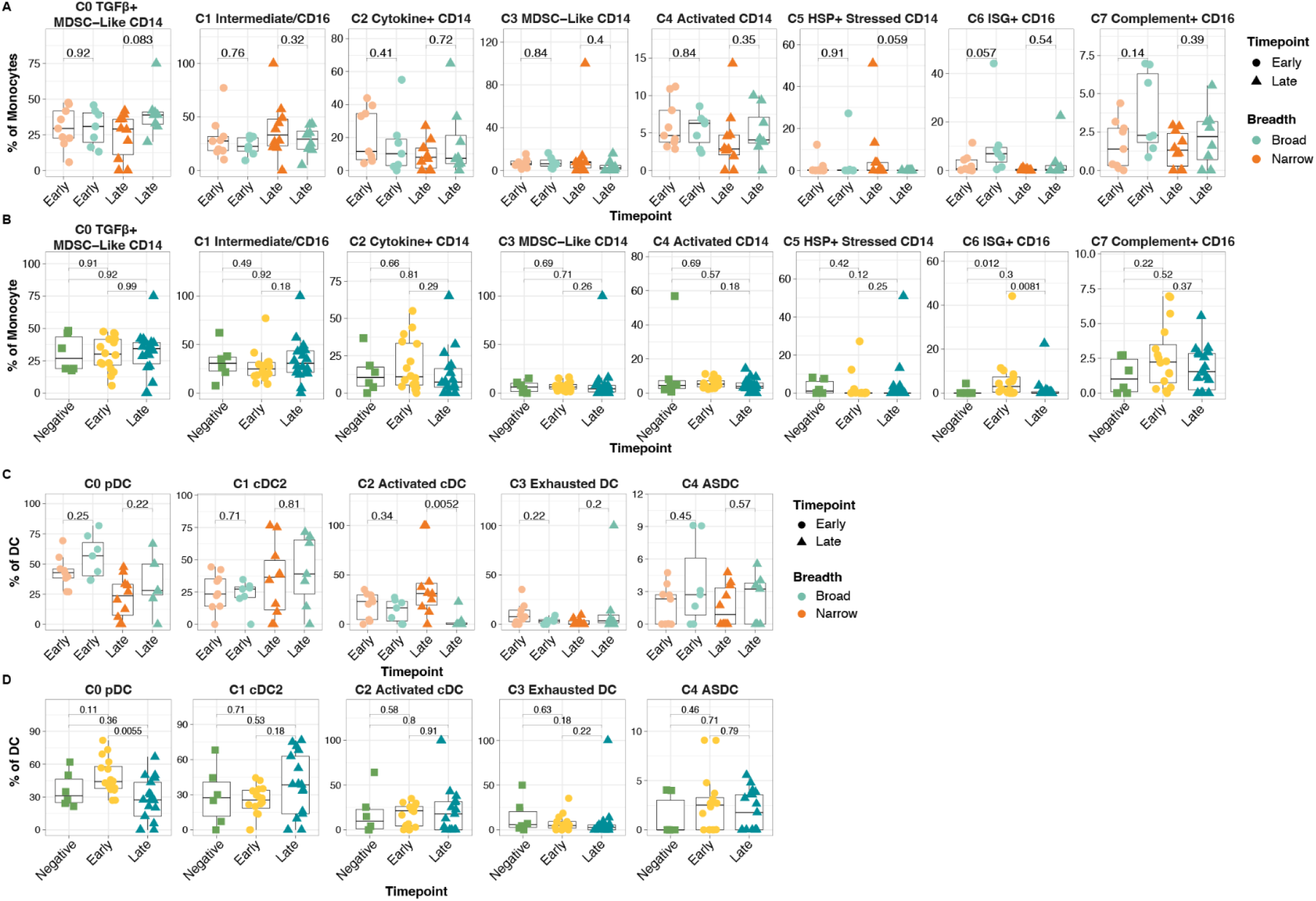
Monocyte and DC transcriptomic clusters. **A.** Boxplots of cluster frequencies amongst monocytes in breadth groups. **B.** Boxplots of cluster frequencies amongst monocytes in infection timepoints. **C.** Boxplots of cluster frequencies amongst dendritic cells in breadth groups. **D.** Boxplots of cluster frequencies amongst dendritic cells in infection timepoints. P-values by two-sided Wilcoxon rank-sum test with Bonferroni’s correction for multiple hypothesis testing. Each point represents one donor.

**Supplementary Fig. 9:**
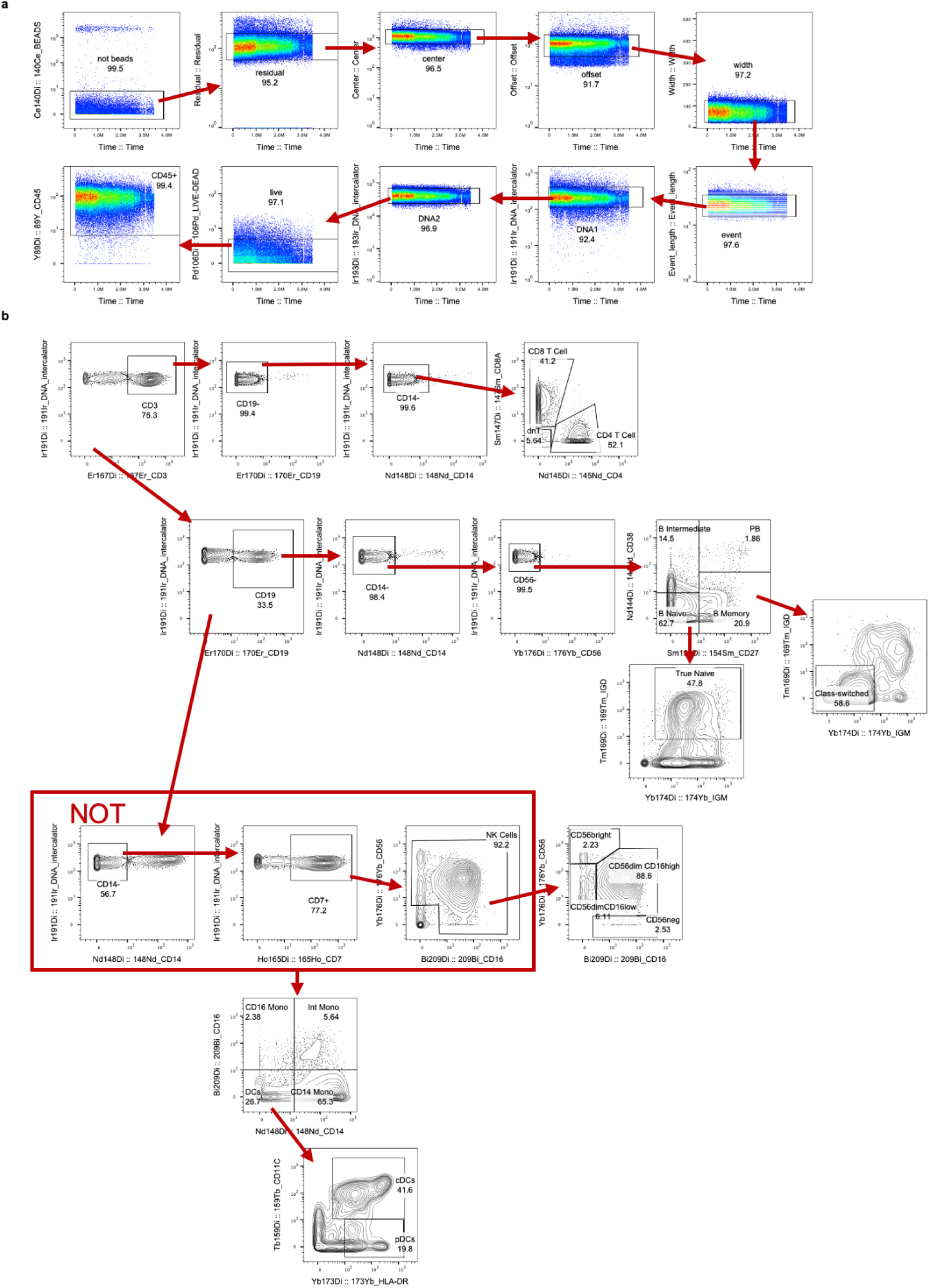
Gating broad cell types in CyTOF data. **A.** Representative flow plots of QC gating following Standard Biotools clean up strategy to identify live, CD45+, singlets. **B.** Representative flow plots of gating used to identify CD4 T, CD8 T, NKs and subsets, Monocytes and subsets, DCs and subsets.

**Supplementary Fig. 10.**
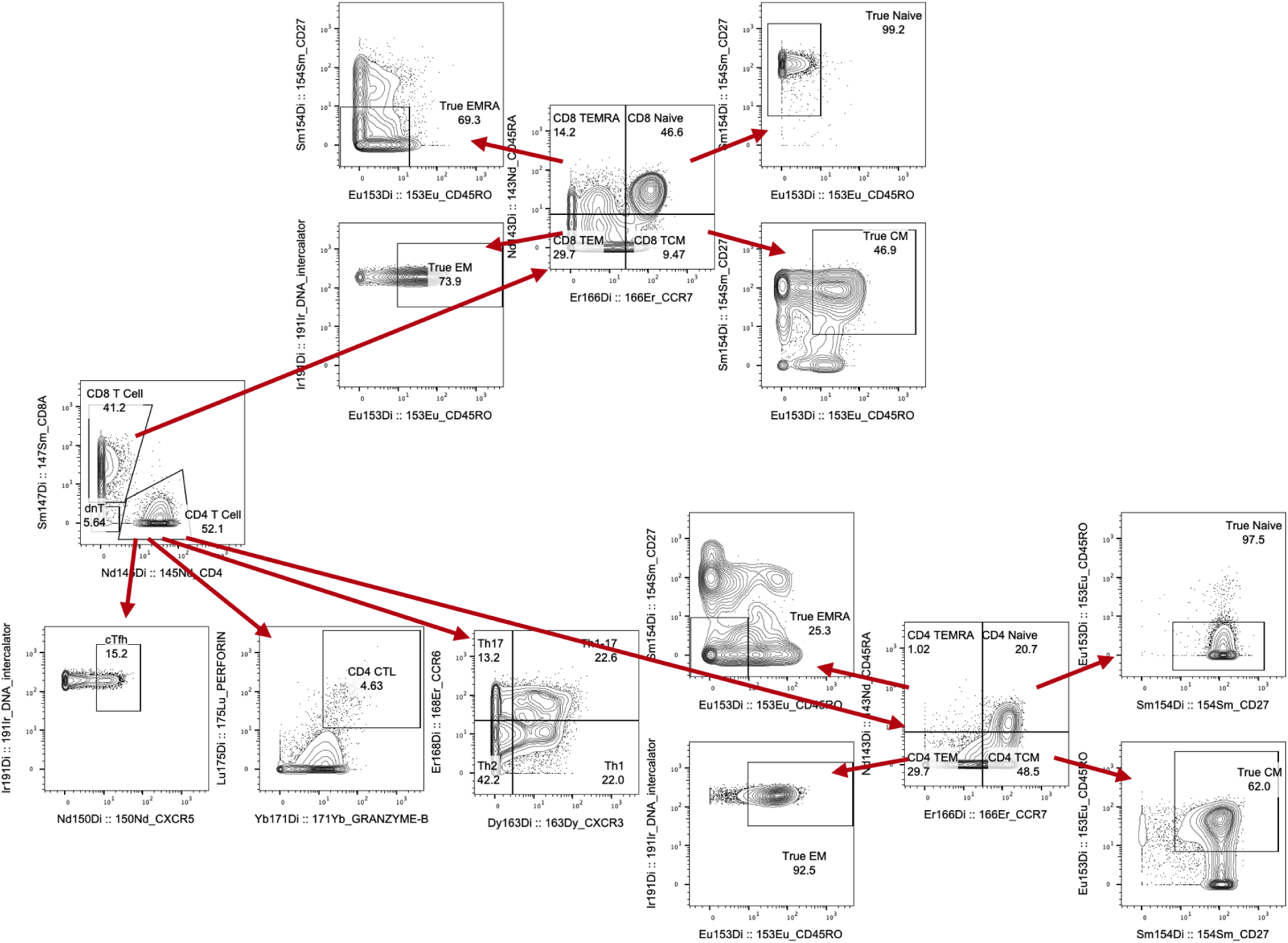
Gating fine T cell subsets in CyTOF data. **A.** Representative flow plots of gating used to identify CD4 T and CD8 T subsets. For CD4 T cells cTfh status, CTL status, helper subtype, and memory subtype was independently identified for each cell.

## Tables

**Table S1: Metadata** Deidentified metadata for each sample.

**Table S2: Titer Data** NT50, PCA, and breadths.

**Table S3: Cell Frequencies** Differences in cell frequencies between timepoints and breadth groups.

**Table S4: Stabl Model Inputs** Feature input for each modality.

**Table S5: Stabl Model Results** Feature selection results for each modality with confounder and donor robustness results.

**Table S6 BTMs** Blood transcriptional modules plotted for each cell type.

**Table S7: Gene Signatures** Previously published gene signatures.

**Table S8: Multinichenet** Top 25 receptor-ligand pairs identified by multinichenet for timepoint and breadth group comparisons.

**Table S9: CyTOF Reagents** CyTOF antibodies and conjugation kits used.

## Author contributions

I.D.K. and C.A.B. conceived the project and designed experiments. I.D.K. acquired single-cell transcriptomic data. I.D.K, U.M., S.D.C, K.P., and M.S., acquired mass cytometry data. I.D.K and C.E.B. performed statistical and computational analyses and generated figures. H.L.I., V.C., R.S.M., K.M., and J.O. obtained patient metadata and processed and stored patient samples used in this study. I.D.K. and C.A.B. wrote the manuscript. All authors reviewed and revised the manuscript.

## Notes

### Competing Interest Statement

The authors have declared no competing interest.

### Summary of Updates

Manuscript was updated. Introduction revised for clarity. Discussion revised for clarity. Figures updated to include experimental model.

